# Improving forest ecosystem functions by optimizing tree species spatial arrangement

**DOI:** 10.1101/2023.10.23.563583

**Authors:** Rémy Beugnon, Georg Albert, Georg Hähn, Sylvia Haider, Stephan Hättenschwiler, Andréa Davrinche, Benjamin Rosenbaum, Benoit Gauzens, Nico Eisenhauer

**Author notes:** Corresponding author: Rémy Beugnon –. Leading author (Rémy Beugnon). Senior author (Nico Eisenhauer). To promote equity between authors, positions were randomized with an exception for first, second, and senior authors.

## Abstract

Reforestation and afforestation programs are promoted as strategies to mitigate rising atmospheric CO_2_ concentrations and enhance ecosystem services. Planting diverse forests is supposed to foster such benefits, but optimal tree planting techniques, especially regarding species spatial arrangement, are underexplored.

Here, using field measurements from the subtropical BEF-China experiment, we simulated leaf litterfall and decomposition, as a function of various spatial arrangements of tree species, from clusters of species to random distributions.

We show that increasing tree species spatial heterogeneity in forests composed of nine tree species led to more evenly distributed litterfall, increased litter decomposition and associated nitrogen cycling by 45%. These effects were amplified with increasing plot species richness, while species functional trait identity and diversity modulated these relationships.

The spatial arrangement of tree species is a critical component determining biodiversity-ecosystem functioning relationships, and considering such spatial aspects is crucial for operationalizing biodiversity-ecosystem functioning theory in realistic re-/afforestation projects.

## Introduction

Promoting carbon sequestration in forests has a strong potential to mitigate increasing atmospheric carbon concentration and thereby climate change^1^. In addition to conserving existing forests, re- and afforestation programs have become an increasingly important practice to counterbalance increasing carbon emissions^1,2^.

The significance of species-rich forests in enhancing and stabilizing forest functioning is particularly relevant in the context of carbon sequestration ^3–6^. Forests of high tree diversity promote productivity and soil carbon storage ^4,7,8^, the latter mostly by enhancing litterfall and litter decomposition, thereby linking aboveground production to soil carbon storage ^9–11^.

Ecosystem functions, such as forest productivity or litter decomposition, are usually assessed at the stand level. Nonetheless, underlying processes (e.g. litterfall, competition for light and nutrients) are spatially constrained ^12–16^, and determined by interactions among neighboring trees ^15,17,18^. For instance, the amount of litter falling on the ground decreases with increasing distance from the individual tree producing the litter, thus its contribution to the litter decomposition ^12–14^. Consequently, the spatial arrangement of tree species defines potential species interactions and ecosystem processes, and thus, underpins biodiversity effects commonly observed at the stand level ^19,20^. Therefore, spatial heterogeneity of tree species should maximize species interactions, and thus, enhance soil carbon sequestration by optimizing litterfall distribution and decomposition ^17,21^. While an increasing number of studies concentrates on examining the effects of spatial heterogeneity on biodiversity-ecosystem functioning relationships at the landscape level (e.g. ^22,23^), there is limited consideration given to the potential benefits of within-habitat spatial heterogeneity of different tree species in enhancing ecosystem functions. Yet, in diverse tree communities where forest management practices might target only one or a few species out of many, the spatial arrangement of trees is a challenging but critical aspect for stakeholders and foresters (^24,25^, https://www.fao.org/3/I7730EN/i7730en.pdf). Therefore, striking the balance between a favorable taxonomic and spatial composition of species-rich forests and a reasonable impact on the feasibility for forest management is a key component of designing future forests that maximize carbon sequestration.

A growing number of international and local initiatives are focusing on finding combinations from the pool of native species for local planting efforts (see Restor: https://restor.eco/). In addition to this search for optimal species combinations, the effects of tree species arrangements on litter dynamics may provide further insight for effective plantation designs.

Beyond species composition, litterfall and decomposition depend much on the functional composition of tree communities, for instance, leaf functional traits such as leaf shape and chemistry ^12,26^. Another consequence of functionally diverse tree communities is a multispecies litter layer with a high litter functional diversity that typically leads to higher decomposition rates ^27,28^. The possibility of combining the characteristics of litterfall and decomposition with the spatial distribution of the litter-producing trees is highly promising for a better understanding and guidance of how re-/afforestation programs can improve forest carbon sequestration and sustainable forest management by informing species selection and arrangement.

Here, we tested the effect of tree species spatial heterogeneity on litterfall and litter decomposition (thereafter “litter dynamics”), and how these processes could be optimized by species diversity and a selection of tree species with appropriate functional traits. First, we simulated nine-species mixture forests following gradients of species spatial heterogeneity, ranging from species planted in blocks to fully random designs including designs with lines or small blocks of species (Fig. 1). Second, using litterfall measurements taken from a large forest Biodiversity-Ecosystem Functioning experiment (BEF-China^29^, Fig. 1, see ^12^), we fitted a species-specific litter distribution model and predicted the litter distribution of each tree previously simulated. Third, using decomposition experimental measurements from the same BEF experiment^12^, we fitted a litter-specific decomposition model and predicted litter decomposition for each square-decimetre on the forest floor of our simulated forests. Thus, we estimated the effect of tree species spatial heterogeneity on litterfall distribution, litter decomposition, and carbon and nitrogen dynamics at the plot level. Fourth, we evaluated how an increasing tree species richness (2, 4, and 9 different tree species) affects the tree species spatial heterogeneity – litter dynamics relationship by replicating the experiment for each stand diversity level. Lastly, we tested the effect of species functional identity and diversity on tree species spatial heterogeneity – litter dynamics relationship.

We expected the spatial heterogeneity of tree species distribution to drive species-specific litterfall spatial patterns, thus litter decomposition processes in nine species mixtures (H1). As interactions among tree species will increase with higher species richness, we expected increasing tree species richness to strengthen the tree species spatial heterogeneity – litter dynamic relationship (H2). Finally, we expected these relationships to be modulated by leaf functional trait identity and diversity (H3, Fig. 1.A).

**Figure 1:**
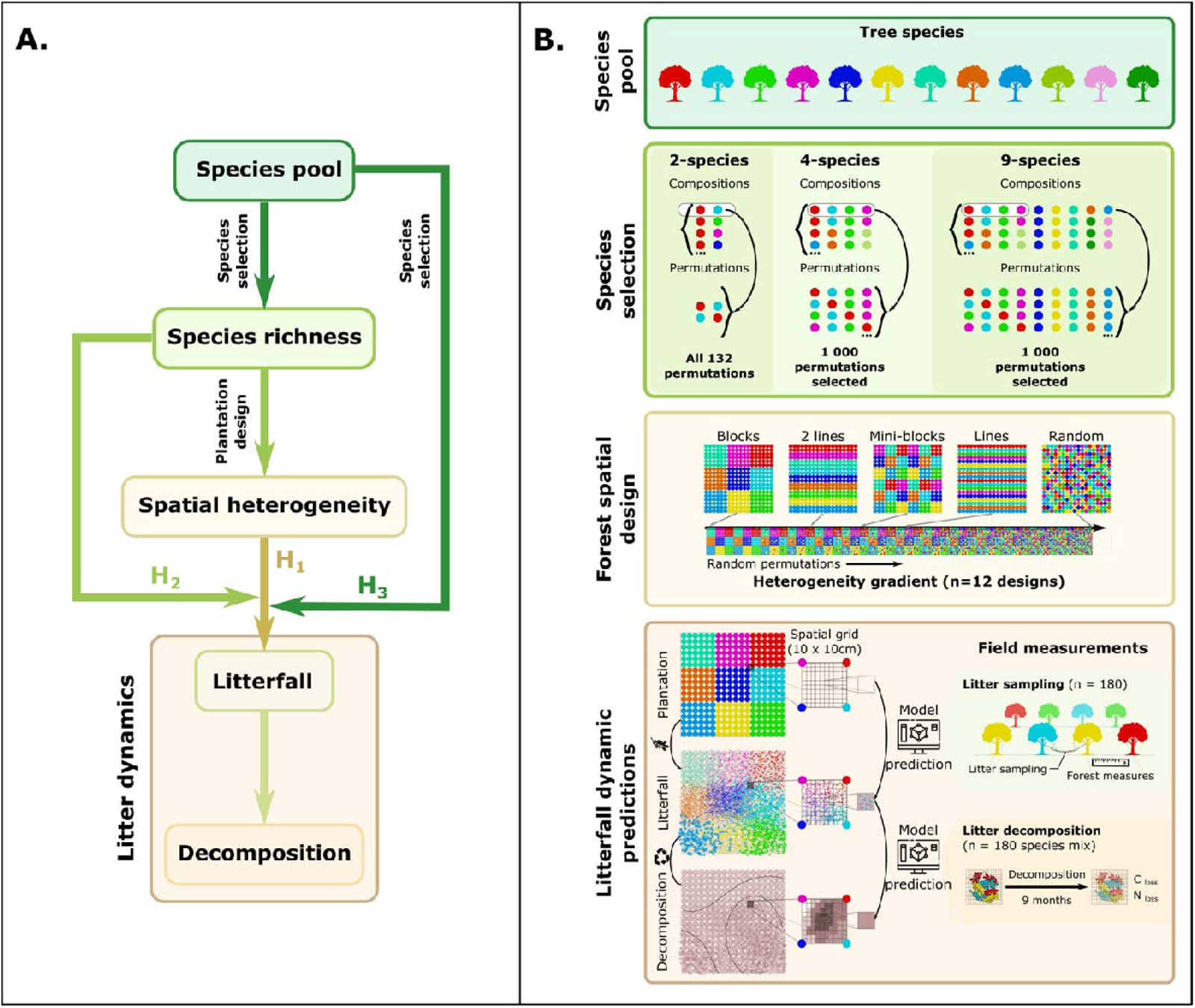
The tested hypotheses (A.) and the associated experimental design (B.). A. Introduction of the hypotheses linking litter dynamics (i.e. litterfall and decomposition) to species spatial heterogeneity (H1) and the mediation by species richness (H2) and species selection (H3). B. Description of the simulation design including species selection, plantation patterns, and the prediction of litterfall dynamics based on field measurements of litterfall and decomposition. Each dot representing a tree is colored according to the species, and each forest stand is composed of 18 by 18 trees.

## Results

### Simulation of forest spatial heterogeneity

To study the effect of tree spatial heterogeneity on forest litter dynamics (litterfall and litter decomposition), we simulated these two processes across a range of plantation designs using inventories, litterfall collections, and leaf litter decomposition field data. In short, we simulated two-, four-, and nine-species mixture forests from a pool of 12 tree species of a Biodiversity-Ecosystem Functioning experiment in subtropical China (BEF-China^29^, Fig. 1). For all the possible mixture combinations, we selected all 132 two-species mixture permutations, 1,000 four-, and 1,000 nine-species mixtures permutations (Fig. 1.B, Suppl. S1, see Methods). Each forest from these 2,132 species mixture permutations was “planted” (i.e. simulated) in forest stands of 18 individuals by 18 individuals of trees planted at a one-meter distance (17 x 17 m forests) with eight different types of spatial distributions from blocks of species to fully random distributions of the species (Suppl. S1). In addition, nine-species mixtures were also planted in smaller blocks, double rows, and single rows of species, to better represent realistic plantation design (Fig. 1.B, Suppl. S1). Taken together, we simulated 21,056 forests (i.e. all species mixture permutations across all planting designs). From tree and species-specific Bayesian models fitted on our experimental litterfall and decomposition data (Fig. 1.B, see Methods and Suppl. S2), we simulated the litterfall distribution of each tree and litter decomposition for each 10 x 10 cm pixel in the simulated forest (Fig. 1.B). We measured tree species spatial heterogeneity as the distance from the null model where nine species are randomly distributed. Planting rows rather than blocks of species increased the tree species spatial heterogeneity, reducing the difference to the random setting by 50% (Suppl S1). However, planting the species in sets of double rows rather than single ones only reduced this spatial heterogeneity difference by 20% (Suppl. S1).

### Tree spatial heterogeneity in forest stands composed of nine tree species reduces litterfall spatial variability and increases litter species diversity

In nine-species mixtures, our simulations showed that the amount of litterfall (average amount of litterfall at forest level) remained constant with increasing tree spatial heterogeneity in the plantation patterns (40.8 ± 5.3 g/dm^2^). However, the variability between pixels decreased with increasing species spatial heterogeneity (from 27.9 to 14.3 g/dm^2^ when comparing block and random designs), showing a homogenization of the litterfall amounts at the forest stand level. Likewise, average litter species richness in a given pixel increased with increasing tree species spatial heterogeneity in nine-species mixture forest stands (from 1.65 to 6.06 species per pixel, Fig. 2A).

**Figure 2:**
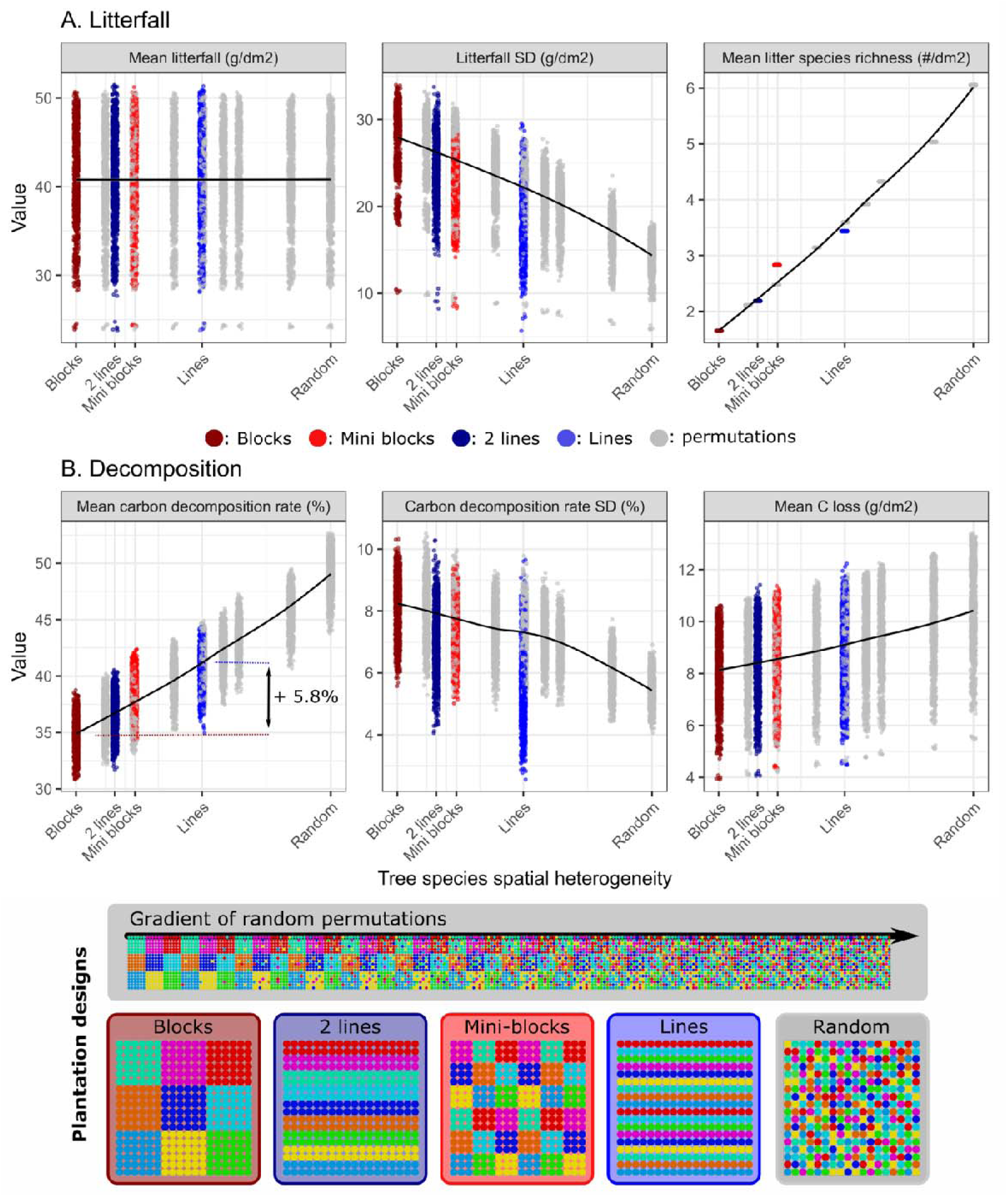
Tree species spatial heterogeneity effect on litterfall (A.) and litter decomposition (B.) in 9-species mixtures. Tree species spatial heterogeneity ranges from blocks of species (dark red) to fully random through the gradient of heterogeneity (gray, Suppl. S1), small blocks of species (3 x 3 trees, light red), double lines (dark blue), and single lines of species (blue). Loess regressions fitted on the tree species spatial heterogeneity gradient (in gray) were added to highlight the patterns.

### Tree spatial heterogeneity increases litter decomposition and reduces the spatial variability of decomposition

Our results for forest stands composed of nine different tree species highlight an increase of decomposition (i.e. carbon loss rate) with increasing tree species spatial heterogeneity (from 34.9 to 49.2% when comparing block and random designs, Fig. 2.B). Similar to the amount of litterfall, we showed a decrease in litter decomposition spatial variability (from 8.2 to 5.5%, Fig 2.B), highlighting a spatial stabilization of the decomposition process with increasing tree species spatial heterogeneity. Changes in litter distribution and its consequences on litter decomposition increased average carbon loss at the forest stand level (from 8.1 to 10.5 g/dm^2^, Fig. 2B). Moreover, planting the nine species in lines rather than in blocks increased the decomposition rate by 5.8%, thus reducing the difference observed between the fully random design and block design by half (e.g. for carbon mean decomposition: block design = 34.9%, line design = 40.7%, random design = 49.2%, Fig. 2.B). Analyses of nitrogen loss showed similar patterns (Suppl. S3), highlighting the stimulating effect of tree species spatial heterogeneity on both, carbon recycling and nitrogen release.

### Tree spatial heterogeneity effect strengthens with tree species richness

In order to evaluate how varying tree species richness modulates the effects of tree spatial heterogeneity on litterfall and decomposition, we repeated our simulations for two- and four-species mixtures and compared those to the results from the nine-species mixtures. We found that the maximum spatial heterogeneity in two- and four-species mixtures introduced with the fully random design reached only the level of heterogeneity from the single rows design in the nine-species mixtures (Fig. 3). We generally observed an increasing strength of tree species spatial heterogeneity effects on the response variables with increasing forest stand tree species richness (Fig. 3A-B). For instance, carbon decomposition rate increased with tree spatial heterogeneity, and this relationship strengthened with increasing species richness (increasing slope from 0.005 to 0.014% when comparing 2- and 9-species mixtures, Fig. 3B). Additionally, when diversity is limited, the influence of tree species spatial heterogeneity on functions such as litter species richness, decomposition rate, or carbon loss plateaus at a low level of heterogeneity. For example, in two-species mixtures, the litter decomposition rate achieves its maximum halfway along the simulated heterogeneity gradient (Fig. 3).

A more detailed examination of the link between decomposition rate and tree species richness revealed a substantial rise in decomposition as species richness increases (from 35 to 40.7% on average when comparing 2- and 9-species mixtures). Similarly, the spatial variability of decomposition rates increased as species richness rose (from 4.33 to 6.98 on average). Consequently, higher tree species richness amplified the spatial variability in litter decomposition, however, this effect became less pronounced in spatially homogeneous planting designs. Conversely, the strength of the biodiversity-ecosystem functioning (BEF) relationship between tree species richness and decomposition rate diminished as spatial heterogeneity decreased. For block designs, these effects become nearly non-significant (e.g. decomposition increases from 33 to 35% between 2- and 9-species mixtures, see Fig. 3C). In summary, our results show that the strength of the BEF relationship is contingent upon the spatial heterogeneity of tree species within the forest.

**Figure 3:**
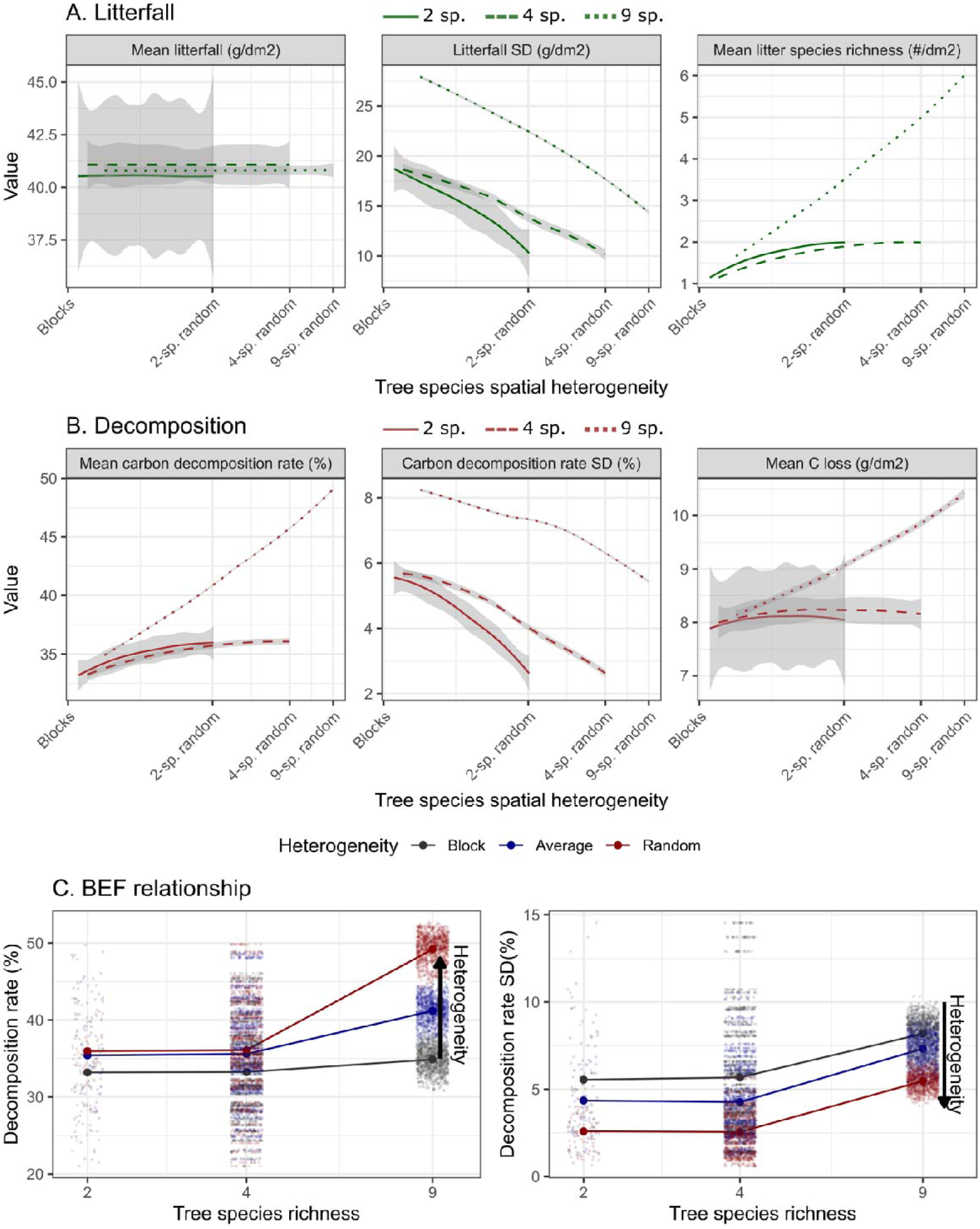
Interaction between tree species richness and tree species spatial heterogeneity on litterfall (A.) and litter decomposition (B.) from two species mixture (solid lines, n = 132) to four (dashed line, n = 1,000) and nine species mixtures (dotted lines, n = 1,000). The gradient of heterogeneity is relative to the 9-species mixture null model (see Methods). C. Biodiversity-Ecosystem Functioning relationship between tree species richness and decomposition across tree spatial heterogeneity level from block to random design (see Suppl. S3 for additional variables).

### Functional determinants of relationships between tree diversity and spatial heterogeneity

We tested the effect of species composition on the relationships between tree species spatial heterogeneity and litterfall dynamics (thereafter heterogeneity-litter dynamics relationship). To derive generalizable recommendations for forest management beyond species identity, we used a functional approach of species and used leaf functional trait (i.e. leaf dry matter content, specific leaf area, and leaf nutrient contents) identity (community weighted mean, CWM) and diversity (community functional richness and dispersion, FRic and FDis respectively). We modeled the intercept and slope of the heterogeneity-litter dynamics relationship as a function of leaf functional trait identity and diversity using a random forest algorithm (see Methods, Fig. 4.A).

On average, our models explained about 15 to 16% of the intercept and slope variability (Fig. 4B). Specifically, our random forest models highlighted leaf functional dispersion (FDis) as the strongest driver of these relationships (Fig. 4C). In addition, both functional axes of the community functional identity (CWM 1 and CWM 2) significantly mediated the heterogeneity-litter dynamics relationship (i.e. variable importance above the 0.25 threshold, Fig. 4C, Suppl. S4). For example, low leaf toughness and high nutrient content (as indicated by high CWM 1) led to an augmentation in decomposition rate (Suppl. S3). Taken together, taking both functional dispersion and identity (i.e. CWMs) into account enables the maximization of both litter dynamics (i.e. effect on intercepts) and heterogeneity-litterfall relationship (i.e. effect on slopes).

**Figure 4:**
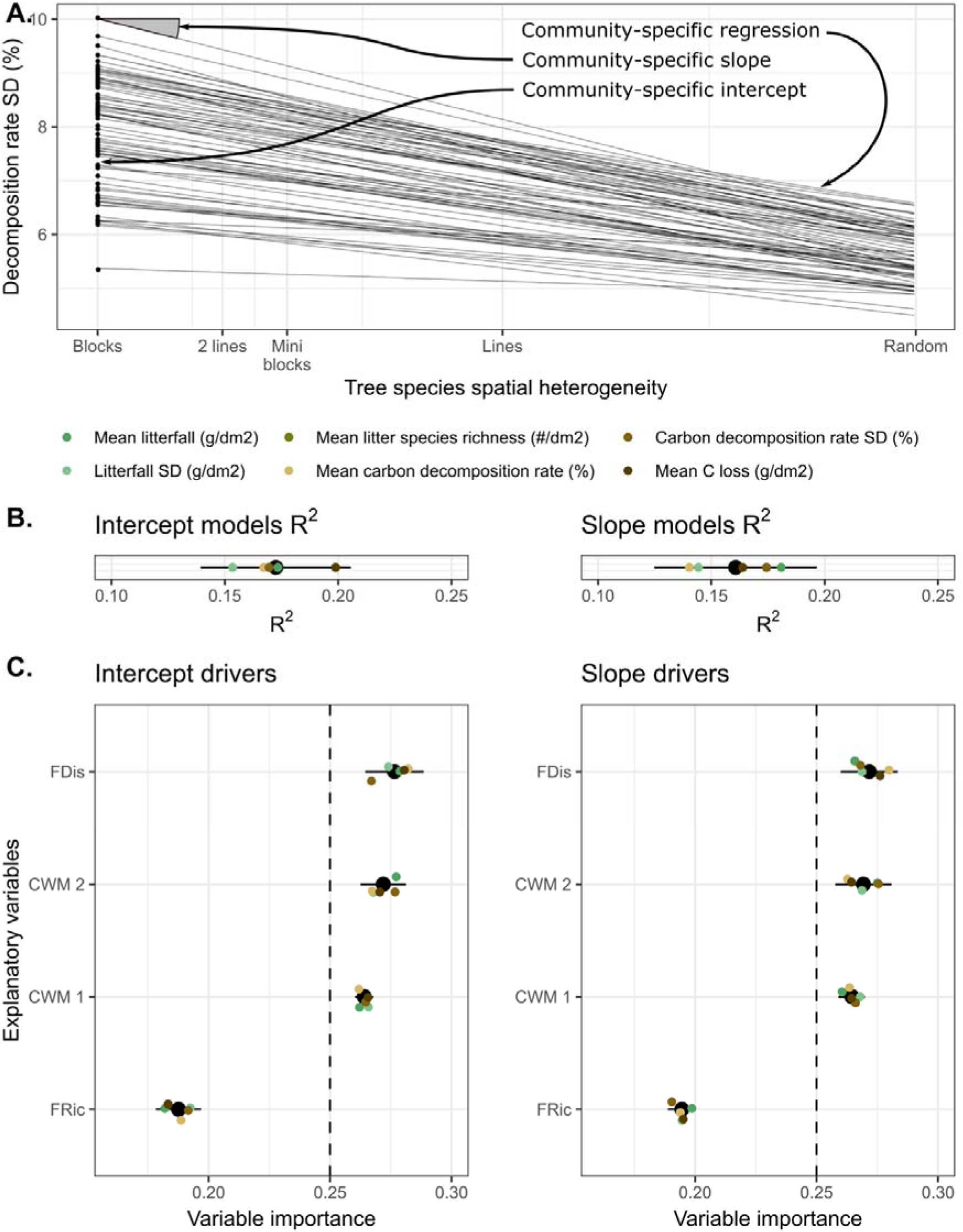
Drivers of the different litterfall and decomposition variables (“Intercept”) and litterfall dynamic – tree species spatial heterogeneity relationships (“Slope”). A. Illustration of the community-specific slope and intercept on decomposition rate spatial variability. B. Explanatory power of the random forest algorithms on the intercept and slope between tree species spatial heterogeneity and each response variable (i.e. “Average litterfall”, “Mean litter species richness”, “Litterfall SD”, “Mean carbon decomposition rate”, “Carbon decomposition rate SD” and “Average C loss”, colored). C. Importance of the leaf functional trait identity (i.e. “CWM 1” and “CWM 2”: leaf functional traits first and second PCA axes, see Method) and diversity (i.e. “FRic”: functional richness, “FDis”: functional dispersion) in the random forest models. Black dots (and lines) represent the mean (and standard error) importance across all variables and colored dots indicate individual variables. The dashed line represents the level below which the variable importance is considered negligible (i.e. 1 / number of variables = 0.25).

## Discussion

By integrating empirical data from extensive field measurements of litterfall and decomposition into simulated forests varying in heterogeneity of the spatial arrangement of tree species, we showed that this heterogeneity strongly affects litterfall and litter decomposition. With increasing spatial heterogeneity, the litter layer is more diverse and most importantly, more evenly distributed across the forest floor without changing the total amount of litterfall. These changes boost decomposition rates and thereby enhance the total amount of carbon processed by the ecosystem (H1). The effects of species spatial heterogeneity are particularly pronounced in species-rich forests (H2) assembled from tree species that are functionally diverse (H3). Our results suggest that the spatial heterogeneity of tree species within forests has a notable impact on the distribution patterns of litterfall and the subsequent litter decomposition, which should be considered in sustainable forest management.

### The effects of species spatial heterogeneity on litterfall and decomposition

Forest tree species spatial heterogeneity has direct effects on litterfall processes. The reduced variability of species-specific litter biomass leads to the spatial homogenization of the litter composition in the forest. The negative correlation between tree spatial heterogeneity and litter spatial variability can be explained by the spatially limited litter distribution (Suppl. S2, ^12,14^). Litter from a particular species tends to stay close to the tree it is shed from, which means that in a block design, different species-specific litters only mix at the edges of the blocks. Conversely, a spatially heterogeneous design aims to overcome this limitation by distributing species more evenly throughout the forest, thereby achieving a more uniform composition of species litter across the entire forest. In addition, processes like wind and topology^30^, but also animal litter translocation can additionally lift the litter spread limitation^31^, therefore reducing the positive effects of tree species spatial heterogeneity on litterfall spatial homogenization.

Our findings suggest that positive effects of tree species spatial heterogeneity on litter decomposition rates (H1) result from an increased litter diversity associated with higher spatial homogenization. Litter species richness was shown to promote litter decomposition at stand-level^12^, for instance, by promoting nutrient transfer between different types of litter^32^. In addition, increasing tree species heterogeneity stabilized litterfall and litter decomposition processes in space. Decomposition is a key ecosystem process linking carbon and nutrient pools between the atmosphere, the biosphere, and the pedosphere, specifically by linking tree primary production and soil carbon stocks^11,33^. Thus, improving and stabilizing litter decomposition by increasing tree species spatial heterogeneity might become a powerful tool to optimize biodiversity benefits for nutrient recycling, carbon sequestration, and climate change mitigation.

### Interactions between tree species diversity and spatial heterogeneity

Numerous studies highlight that high species diversity enhances multiple forest ecosystem functions^34^, including those directly relevant for forest management, namely forest productivity and carbon sequestration ^4,8,35,36^ and the stability of ecosystem functions^6^. Our findings add the tree species spatial heterogeneity as an additional, and so far largely neglected, component to how tree species diversity can influence ecosystem functioning. This positive relationship between tree species richness and decomposition as an important ecosystem function is maximized when the different tree species are planted fully randomly and remains limited when the different tree species are aggregated into blocks, i.e. patches of single species (Fig. 3C). The interaction between tree species richness and tree species spatial heterogeneity highlights the crucial aspect of species interactions for ecosystem functioning^17,18^. In addition, the modulating impact of tree species spatial heterogeneity on the tree species richness effect may be a key to the understanding of the variability in the strength of BEF relationships in the literature ^37,38^. For instance, experimental designs typically differ substantially, specifically between completely random designs (e.g. BEF-China^29^) to more clustered designs (e.g. FORBIO experiment^39^), and a single experiment does not commonly include variable spatial heterogeneity in addition to species diversity. Particularly in situations with limited species diversity, our findings indicate that the influence of tree species richness is minimal and falls behind the impact of species spatial heterogeneity. Consequently, as species richness decreases, prioritizing the consideration of tree species spatial heterogeneity becomes increasingly decisive for enhancing ecosystem functioning.

Taken together, these results highlight the need to integrate small-scale spatial distribution of species into BEF analyses for a better understanding of processes with strong spatial constraints. Here, the significance of heterogeneity in biodiversity-ecosystem functioning (BEF) relationships appears to be closely tied to the scale at which it is examined^13^. It necessitates not only an awareness of its spatial extent but also a comprehension of the fundamental processes at play. In particular, processes such as decomposition exhibit a strong connection to the dispersal capabilities of litter and decomposer communities. Consequently, defining the appropriate spatial scale of interest ranges from just a few centimeters for nutrient transfers between different litter types to a few meters when considering the spread of microbial decomposers (Bahram et al., 2016).

### Functional trait mediation of tree species spatial heterogeneity effects

The positive effects of forest spatial heterogeneity are not only affected by species richness but also depend on the functional composition of the forest community. Our findings show that functional diversity is a particularly strong determinant of the positive heterogeneity effects (H3). This confirms findings from previous studies that highlight the central role of functional diversity in decomposition processes^40–42^. Optimizing litter dynamic processes by promoting forest diversity and species spatial heterogeneity, therefore, aligns with the repeated calls for enhancing functional diversity in re- and afforestation programs^5^.

### Consequences for forest management

When it comes down to the reality of forest management decisions, species-rich forests are often considered too costly and even as not feasible practically, despite the overall beneficial effects of species diversity on forest sustainability (e.g., productivity, stability, nutrient cycling). Our model simulations specifically addressed the question of how different planting designs affect decomposition-related ecosystem processes to help finding a compromise between the beneficial tree diversity effects and the feasibility of management. We demonstrated that the easiest management solution with a simple block design ranks lowest in terms of decomposition-related processes. On the other hand, a completely random planting design, maximizing spatial heterogeneity is the ideal scenario from an ecological perspective, but rendering management difficult. An interesting compromise could be planting trees in lines that can still strongly improve forest functioning compared to block designs while keeping forest management feasible. Enhancing spatial forest heterogeneity for improved and sustainable forest ecosystem functioning, however, has to be considered in comparison to other options previously suggested as contributing to sustainable forest management. Additionally, we showed that tree functional traits could explain variability in litterfall and litter decomposition rates and their relationship to species spatial heterogeneity. Thus, trait-based informed management might allow the optimization of forest functioning and provide general guidelines for managers by relaxing the constraints on our mechanistic understanding of these processes, imposed by species identity^43,44^.

## Conclusions

Climate change and biodiversity loss are listed as major global change drivers of ecosystem functioning^45,46^. Re- and afforestation are identified as nature-based solutions to mitigate climate change by counteracting the rising atmospheric CO2 concentration and safeguarding ecosystem functioning ^1,2,47^, being especially effective when considering tree species-rich plantations ^5,47^. However, the diversification of existing and newly planted forests may be challenging for current management practices, requiring different options for optimal solutions. Our findings provide a clear quantification of different planting solutions for key ecosystem processes, demonstrating that any promotion of species spatial heterogeneity enhances the biodiversity effect on ecosystem functioning. Simply transforming plantations of blocked species clusters into adjacent rows of different species was shown to already improve ecosystem functioning considerably, e.g. by increasing decomposition by 45%, while having acceptable consequences for forest management. To develop sustainable management strategies, future research should integrate both fundamental and applied perspectives in experimental designs, with a specific focus on studying species interactions and their ecosystem consequences across spatial scales.

## Acknowledgments

This work was supported by the Deutsche Forschungsgemeinschaft (DFG, German Research Foundation – 319936945/GRK2324; 452861007/FOR5281). We gratefully acknowledge the support by the German Centre for Integrative Biodiversity Research (iDiv) funded by the German Research Foundation (DFG– FZT 118, 202548816). We thank the TreeDì and Experimental Interaction Ecology research groups for their support, especially Georg Hähn and the many local helpers for their help with the field sampling. N.E. acknowledges funding by the DFG (Ei 862/29-1, Ei 862/31-1).

## Data availability

Data will be freely available and uploaded on DRYAD

## Code availability

Code will be freely available on Zenodo (attached to the submission for reviewers)

## Authors’ contributions

RB, NE, StH: designed the study; RB, GH, AD, SyH: collected field data; RB, GA, BR, BG: designed the simulations; RB, GA, GH: analyzed model outputs; RB, GA: prepared the first draft, RB, GA, GH, AD, SyH, BR, BG, NE, StH: revised the manuscript.

