## Supplementary material for "Improving forest ecosystem functions by optimizing tree species spatial arrangement": Methods

### Study site

The study site is located in southeast China near Xingangshan city (Jiangxi Province, 29.08–29.11° N, 117.90–117.93° E). Our experimental site is part of the BEF-China experiment (site A ^1^), and it was planted in 2009 after a clearcut of the previous commercial plantation. The region is characterized by a subtropical climate with warm, rainy summers and cool, dry winters with a mean temperature of 16.7°C and a mean annual rainfall of 1821 mm ^2^. Soils in the region are Cambisols and Cambisol derivatives, with Regosol on ridges and crests^3,4^. The natural vegetation consists of species-rich broad-leaved forests dominated by *Cyclobalanopsis glauca*, *Castanopsis eyrei*, *Daphniphyllum oldhamii*, and *Lithocarpus glaber*^1,5^.

### Field sampling design

To identify the effect of tree spatial organization on litterfall distribution and decomposition, we measured litterfall and decomposition between 180 pairs of trees with varying neighborhood composition and species richness. Each pair consisted of two trees next to each other (1.28 m), and we defined its neighborhood as the ten trees directly adjacent in the planting grid^6^. Each unique species pair was replicated three times at every tree species richness levels (plot species richness of 1, 2, 4, 8, and ≥ 16 species) whenever possible (“broken stick” design^1^). In total, we surveyed 24 unique combinations of tree species resulting in a total of 180 pairs of trees in 52 plots^7^.

### Tree biomass estimations

Tree biomass, used for fitting litterfall models, was predicted for all tree pairs and their neighbors using tree basal area (BA) and species-specific allometric relationships estimated on the pair of trees. (1) Circumference at breast height (CBH) was measured in September 2018 for all pairs of trees and their direct neighbors in order to calculate the basal area of these trees as 𝐵𝐴 = (𝐶𝐵𝐻)^2^/4𝜋. (2) Tree height was measured for the pair of trees, and tree biomass was calculated following Huang et al. (2018)^8^. Tree pairs’ BA and biomass were used to estimate species-specific allometric BA-biomass relationships and predict the tree biomass for all neighboring trees^9^.

**Leaf functional traits measurements**

Leaf functional traits were assessed at the species- and plot-level in September 2018, following Davrinche and Haider (2021)^10^. For each species in each plot, several leaf samples were collected, and the reflectance spectra were measured using ASD FieldSpec® 4 Wide- Resolution Spectroradiometer (Malvern Panalytical Ltd., Malvern, United Kingdom). Leaf functional traits were predicted from the reflectance spectra of a calibration dataset of the same species, where both reflectance spectra and leaf functional traits were measured. For leaf morphological traits – specific leaf area (SLA, leaf area divided by dry weight) and leaf dry matter content (LDMC, ratio of leaf dry mass to fresh mass) – fresh and dry weights were measured before and after drying for 72 h at 80°C. To obtain SLA, leaf areas were measured from scans with a resolution of 300 dpi of the fresh leaves using the WinFOLIA software (Regent Instruments, Quebec, Canada). Leaf chemical contents (carbon: C, nitrogen: N, phosphorus: P) were measured from dried leaves ground into a fine powder (Mixer Mill 400, Retsch, Haan, Germany). About 5 mg of leaf powder was used to determine C and N content with an elemental analyzer (Vario EL Cube, Elementar, Langenselbold, Germany). A 200 mg subsample was used to measure P content via nitric acid digestion and spectrophotometry using the acid molybdate method. The filtrate resulting from nitric acid digestion was analyzed with atomic absorption spectrometry (ContrAA 300 AAS, Analytik Jena, Jena, Germany) for magnesium (Mg), calcium (Ca) and potassium (K) content. The relation between the leaf spectra of the calibration samples and the leaf traits was analyzed in the software Unscrambler X (version 10.1, CAMO Analytics, Oslo, Norway) to predict species- and plot-specific leaf functional traits^10^.

### Litterfall sampling

In September 2018, a litter trap of 1 m² was set up at a height of 1 m above the soil surface between each pair of trees^7^. Litter was collected during December 2018 to cover the main litterfall season in the region^11^. To measure litterfall composition, each leaf of the litter trap was sorted and identified to species level. Each species' litter was dried at 40°C for two days and weighed (± 0.1 g).

### Decomposition measurements

We performed a decomposition experiment between the pair of trees to measure

total leaf litter decomposition. Large-mesh litter bags (10 cm x 10 cm) were built using a 5 mm-mesh for the upper part of the bag to provide access to macro-decomposers, and a 0.054 mm-mesh at the bottom to prevent loss of fine leaf litter particles, and filled with 2 g (± 0.01 g) of dried litter according to litter trap species composition (i.e., species-specific biomasses) of the different pairs of trees. Therefore, the litter composition of the litterbags matched exactly the litter composition (i.e., species-specific litter masses) collected in the litter traps of the corresponding pair of trees. The litterbags were installed in December 2018 and covered by a 1 m x 1 m grid to prevent dislocation by heavy rainfalls (1 cm mesh size). In September 2019, i.e., after nine months of decomposition and before the start of litterfall, litterbags were collected, water-cleaned and dried at 40°C for two days. The residual litter was weighed (± 0.01 g) and milled.

Litter C and N content after decomposition were measured from the residual litter with an elemental analyzer (Vario EL Cube, Elementar, Langenselbold, Germany) and corrected for soil contamination following Beugnon, Eisenhauer et al. (2023)^7^. C and N loss rates (%) from the litterbags during the period from December 2018 to September 2019 were calculated.

### Simulations

All simulations, statistical analyses, and data visualizations were performed using R software version 4.3 (R Core Team^12^), and R-scripts are provided to the readers on our Zenodo directory (<https://github.com/remybeugnon/Beugnon-et-al-2023-heterogeneity-litter-dynamics>). To study the effect of tree spatial heterogeneity on litterfall and litter decomposition, we simulated a range of forest designs and predicted litterfall and litter decomposition using Bayesian models fitted to our field data for 2-, 4- and 9-species mixtures (Fig. 1), to ensure a symmetric partitioning of the block designs (see Suppl. S1).

#### Species selection

From the 24 species planted in BEF-China experimental Site A, 12 species were measured during the litterfall sampling and used in the decomposition experiment^7^. From all the possible 2-, 4-, and 9-species mixtures combinations, we selected all 132 2-species mixture permutations (e.g. Sp1-Sp2, Sp2-Sp1 …), and randomly chose 1,000 4- and 1,000 9-species mixtures permutations, totaling 2,132 species mixtures permutations. Permutations of the species arrangement for a given species mixture allow for randomized species interactions (see example in Suppl. S1). Those species mixture permutations were then “planted” (i.e. simulated) across different planting designs. Tree biomass was fixed to the average species-specific biomass measured in the experiment to avoid confounding effects of species interaction on biomass^13^ and heterogeneity treatment in the simulations. Therefore, our experiment will not reflect the positive species richness – productivity relationship ^8,9^.

#### Plantation designs

The forests were planted in an 18 by 18 grid of trees following the species selection within a plot area of 289 m^2^ (i.e. 11,211 individual trees per hectare). Tree species spatial heterogeneity was manipulated by using different plantation designs, with a block design - meaning that all individuals from a given species were planted together (e.g. 36 individuals per species in each corner of a plot for a four-species mixture) - and a fully random spatial arrangement of the species at the two extremes of the spatial heterogeneity gradient (Fig. 1, Suppl. S1). To create a gradient of spatial heterogeneity, trees were randomly permutated, and six designs were selected to create a regularly homogeneously distributed gradient from the block to the random design (Suppl. S1). In an attempt to closer connect planting design options to a realistic forest management context, we further added specific spatial designs for 9-species mixtures: mini-blocks (i.e. smaller blocks of only 3x3 individuals per species), double-lines, and single-lines (i.e. one single line of planted individuals per species, Suppl. S1). Tree species spatial heterogeneity was measured as the distance from the random distribution of species in terms of average heterospecific direct neighbours. For each plantation design, we calculated litterfall and decomposition rates based on models fitted on empirical data.

### Litterfall and decomposition predictions

*Litterfall model*

To predict the litterfall within our simulated forests, we fitted litterfall models on empirical data, assuming that the leaf litterfall mass L_i_ of species i at any place within the forest is a function of the tree aboveground biomass and of the distance of the surrounding trees of the same species. We predicted litter biomass L_i_ accordingly as:

$L_{i}=b_{1i}\sum_{j=1\ldots n} B_{ij}+b_{2i}\sum_{j=1\ldots n} d_{ij}^{-1}+b_{3i}\sum_{j=1\ldots n} B_{ij}d_{ij}^{-1}$	(1)

where b_1i_, b_2i_ and b_3i_ are species-specific coefficients capturing the influence of (1) the biomass of the n surrounding trees of species i, B_ij_, (2) the distance to n trees of species i, d_ij_, and (3) the interactive effect of biomass and distance. For the model, we considered the 12 trees surrounding the litter traps (i.e., $n\leq12$; see Suppl S4).

*Decomposition model*

We fitted litter decomposition models using the diversity-interaction modeling framework^14^. The model distinguishes between species identity and species diversity effects on litter decomposition. Because any potential non-additive litter diversity effects are difficult to predict, we assumed that diversity effects were purely additive. Hence, litter decomposition was described as:

$D=\sum_{i} \beta_{i}P_{i}+\sum_{i,j} (\alpha_{i}+\alpha_{j})P_{i}P_{j}+b_{4}L+b_{5}S$	(2)

where β_i_ and α_i_ are species-specific coefficients capturing identity and diversity effects, respectively, with P_i_ being the proportion of litter mass from species i in the total litter mixture mass L. The coefficients b_4_ and b_5_ capture the effects of total litter mixture mass, L, and litter species richness, S.

The litterfall and decomposition models were fitted with Rstan ^15^, using four Markov chains and 3,000 iterations (1,000 as warm-up). Model fit, posterior distribution and quality indices are provided in Suppl. S2.

*Predicting litterfall and decomposition in simulated forests*

Using the mean coefficients retrieved from our litterfall model, we predict litter mass for every species at any given 10 x 10 cm pixel within our simulated forests. This allowed us to determine species-specific litter mass and species richness for each pixel. This was then used to predict decomposition rates using the mean coefficients from the decomposition model.

### Leaf functional trait effects on litterfall and litter decomposition - tree species heterogeneity relationships

To evaluate the potential effects of leaf functional traits on the relationship between tree species spatial heterogeneity and litterfall and litter decomposition, we used random forest models. For each heterogeneity ~ litter variable (i.e. average litterfall, litterfall spatial variability, litter species richness, average decomposition rate, decomposition rate spatial variability, average carbon loss) relationship, we tested how much variation in intercept and slope can be explained by community-weighted means of the leaf economic spectrum axes (i.e. the first two PCA axes of the different leaf functional traits; see Suppl. S3), and the functional diversity (functional richness, functional dispersion) of the forest species mixture simulated. For calculating parameters and performing random forest analyses, we used the FD^16^, FactoMineR^17^ and randomForest^18^ R packages.
