## Supplementary material for "Improving forest ecosystem functions by optimizing tree species spatial arrangement": Suppl. S1

### Suppl. plot designs

Beugnon et al.

#### Number of species per plot

We constructed plots featuring mixtures of 2, 4, and 9 different species to create evenly distributed groups of species within 18 by 18 tree plots. To elaborate, we established two sets of 18 by 9 tree blocks for 2-species mixtures, four sets of 9 by 9 tree blocks for 4-species mixtures, and nine sets of 6 by 6 tree blocks for 9-species mixtures. Additionally, the 18 by 18 tree plots enabled us to design smaller 3 by 3 tree blocks within 9-species mixtures.

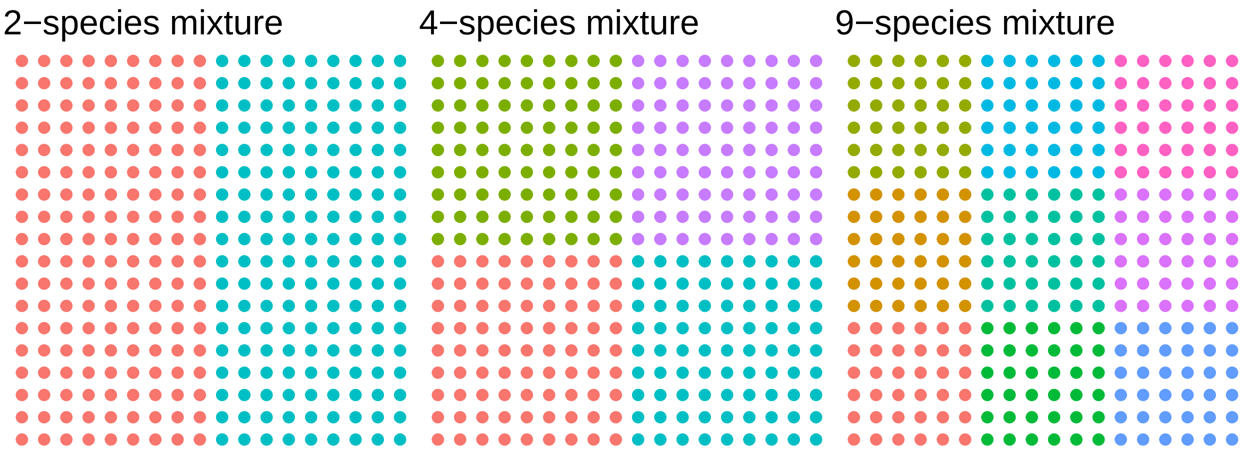

### Permutation gradient

#### 9-species mixtures

Black lines highlight selected permutations

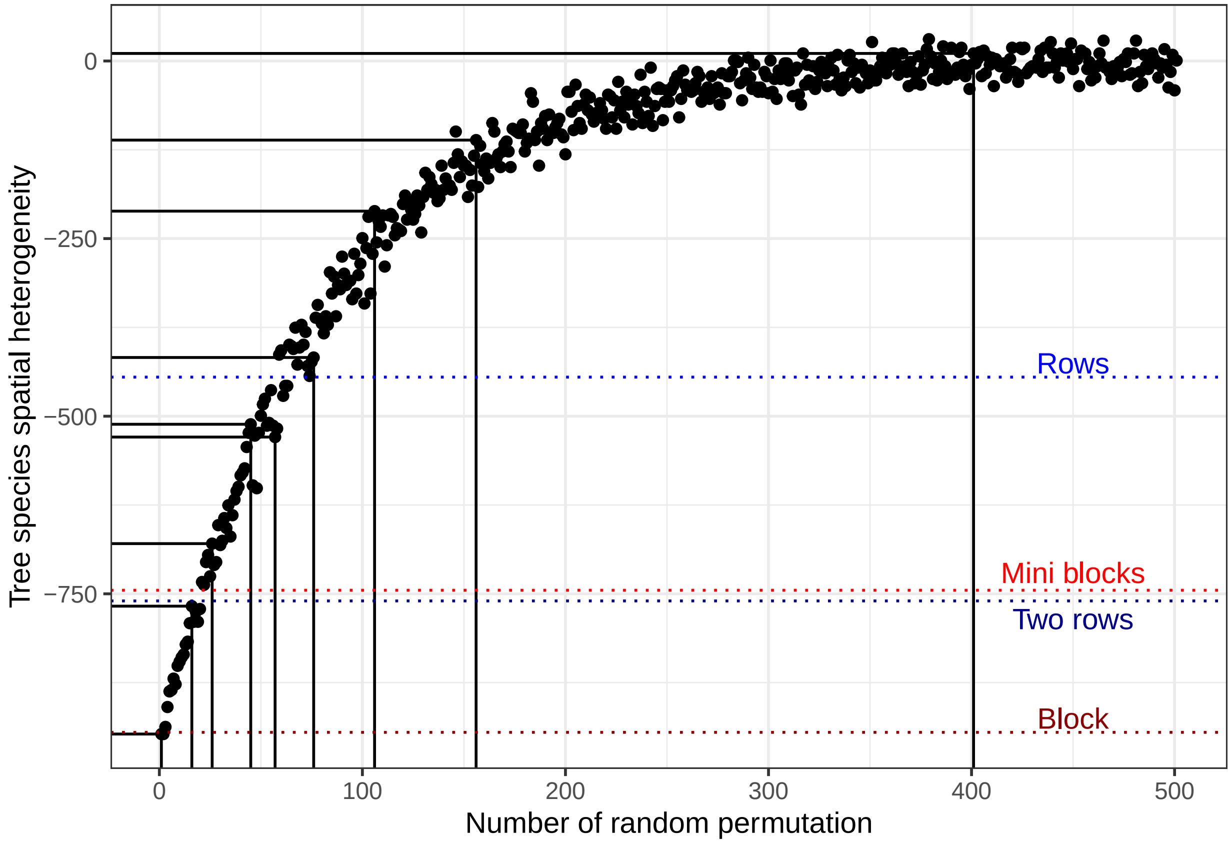

**4-species mixtures**

Black lines highlight selected permutations

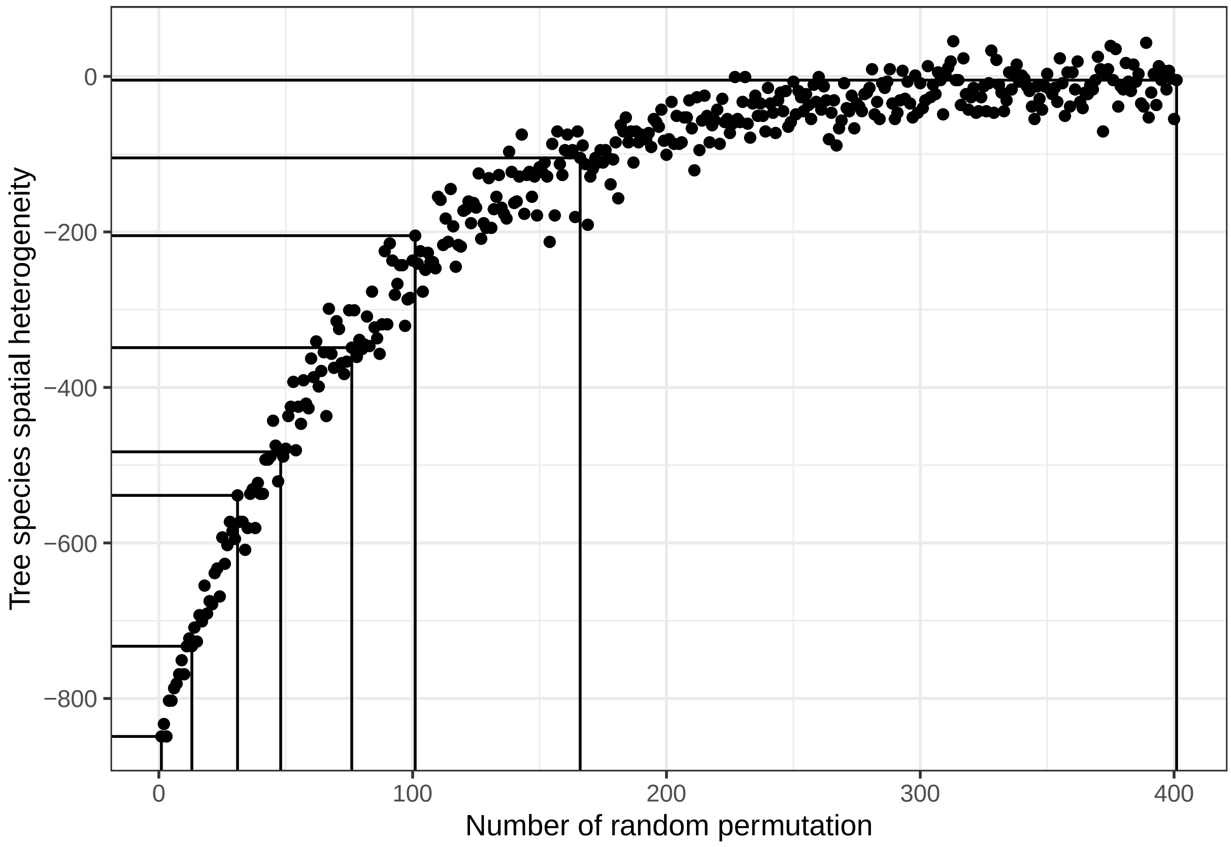

#### 2-species mixtures

Black lines highlight selected permutations

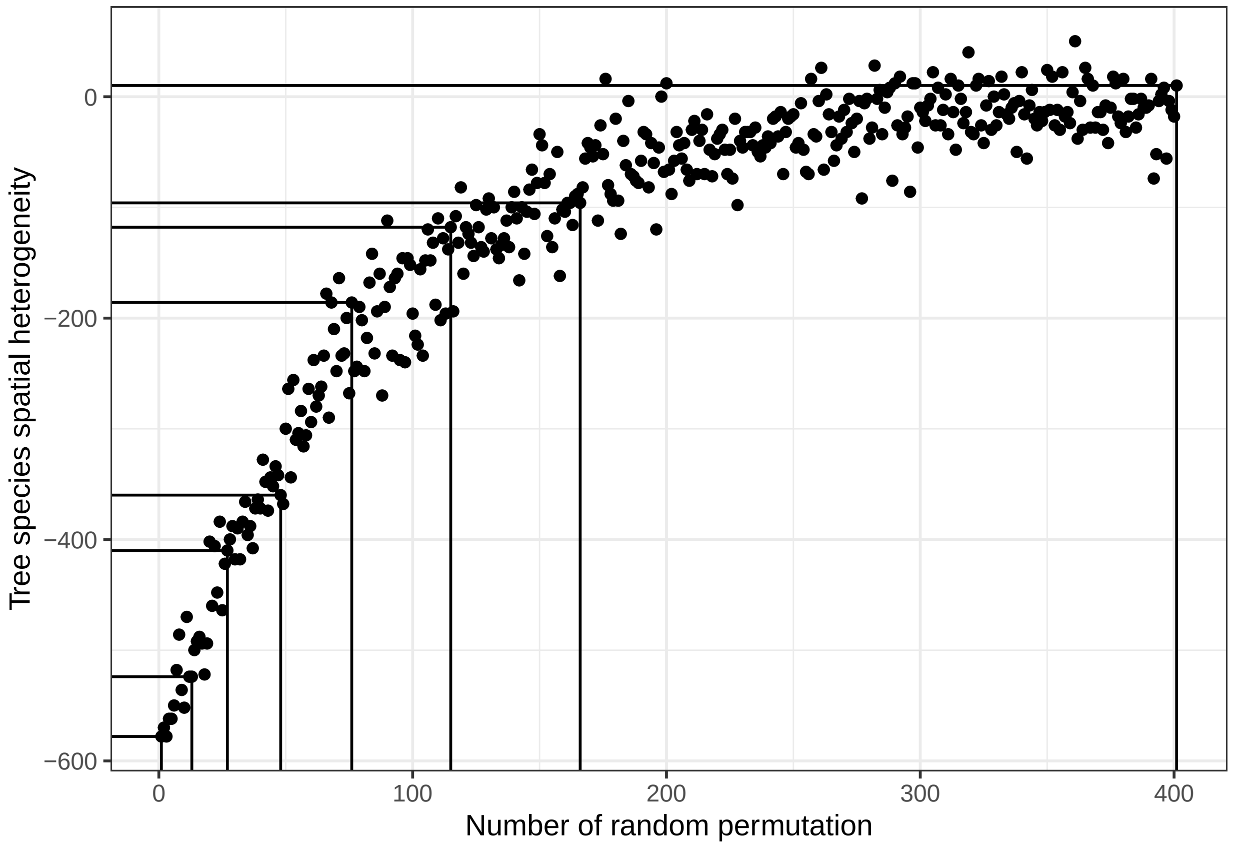

#### Plot designs

Plot spatial designs selected for the study. Colored dots represent the species distributed on the 18 by 18 tree forests

##### 9-species mixtures

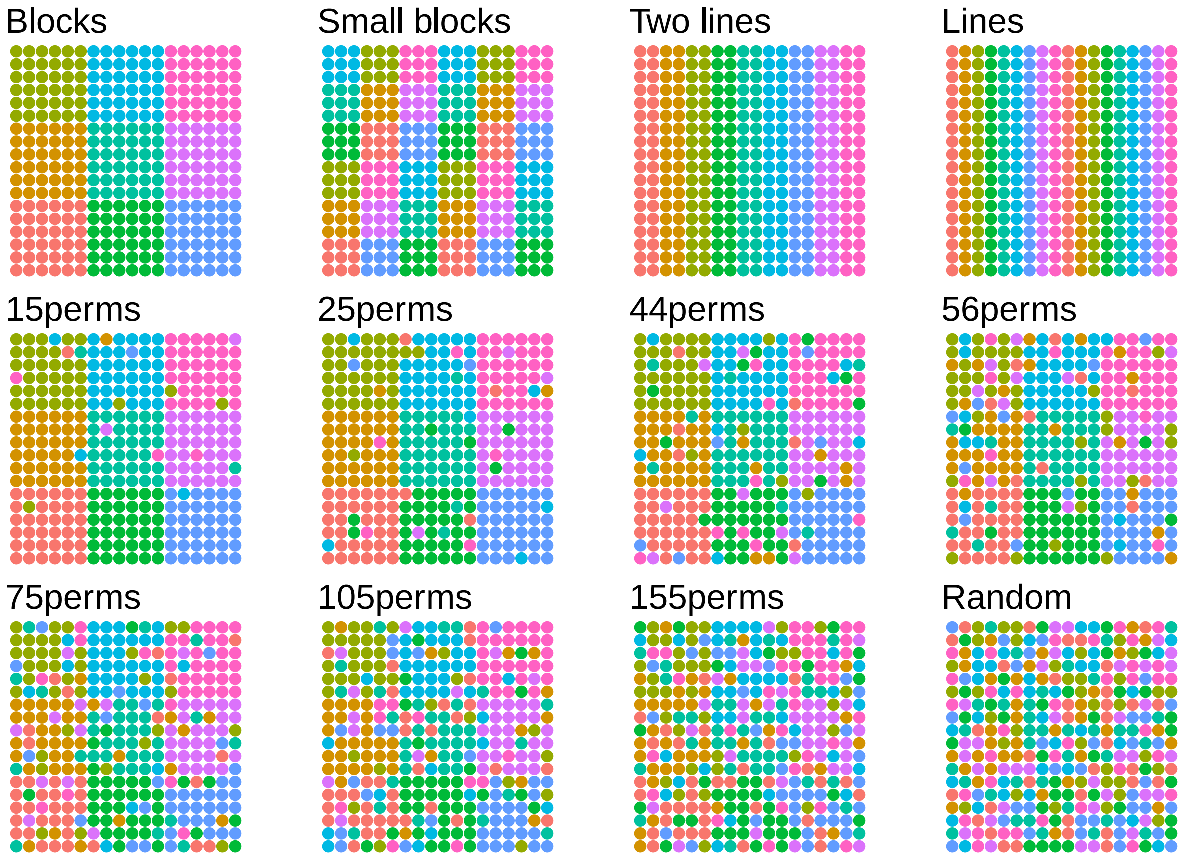

**4-species mixtures**

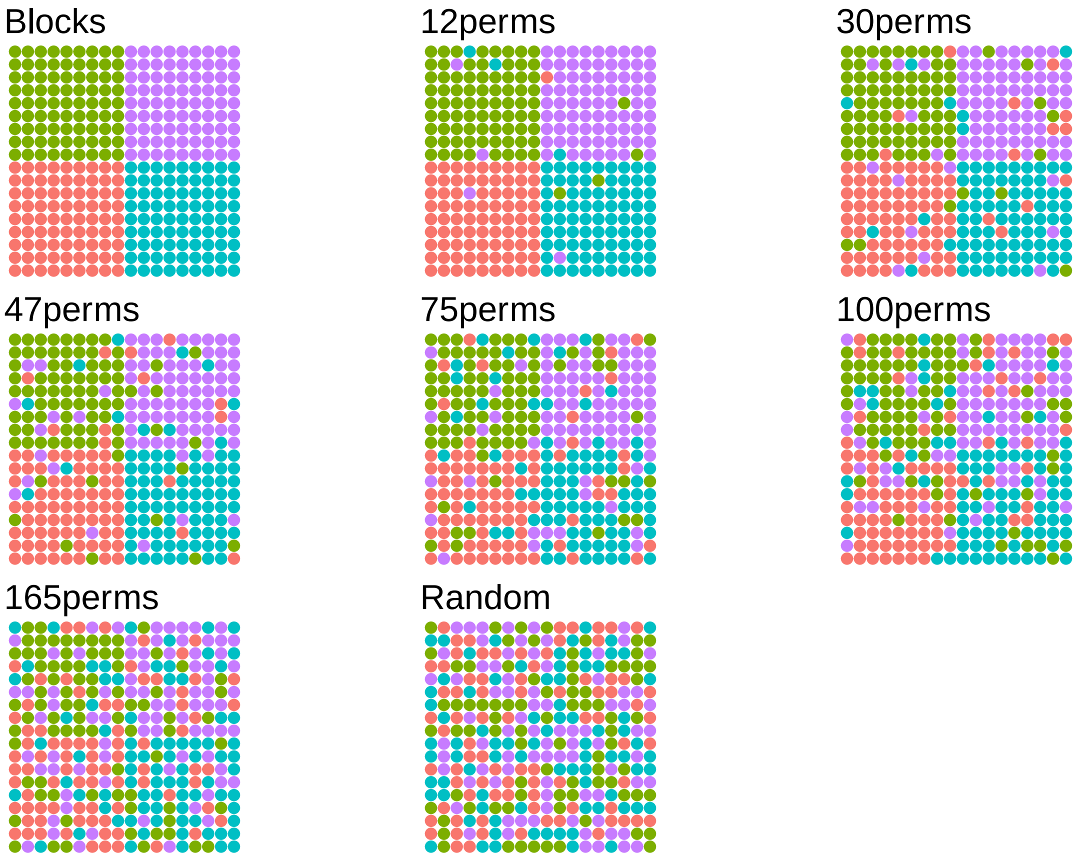

**2-species mixtures**

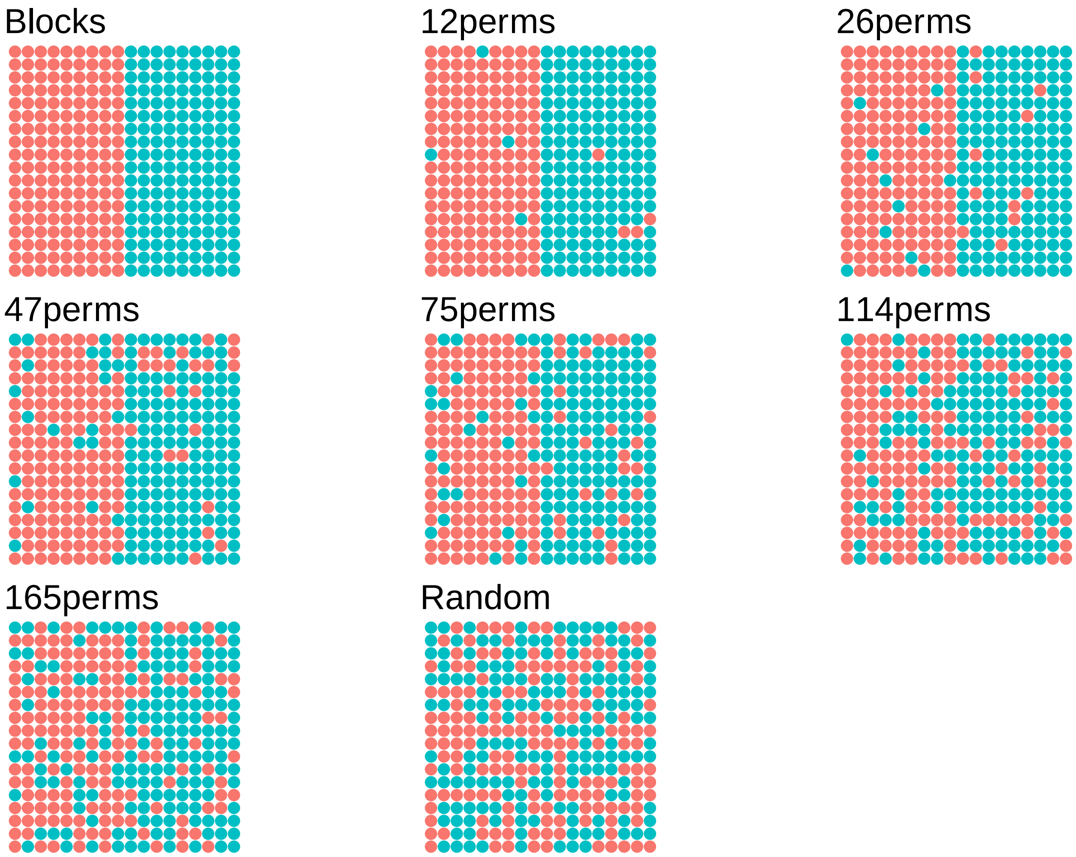
