## Supplementary material for "Improving forest ecosystem functions by optimizing tree species spatial arrangement": Suppl. S2

### Supplementary material S2

#### Litterfall distribution models

##### Datasets

###### Input variables

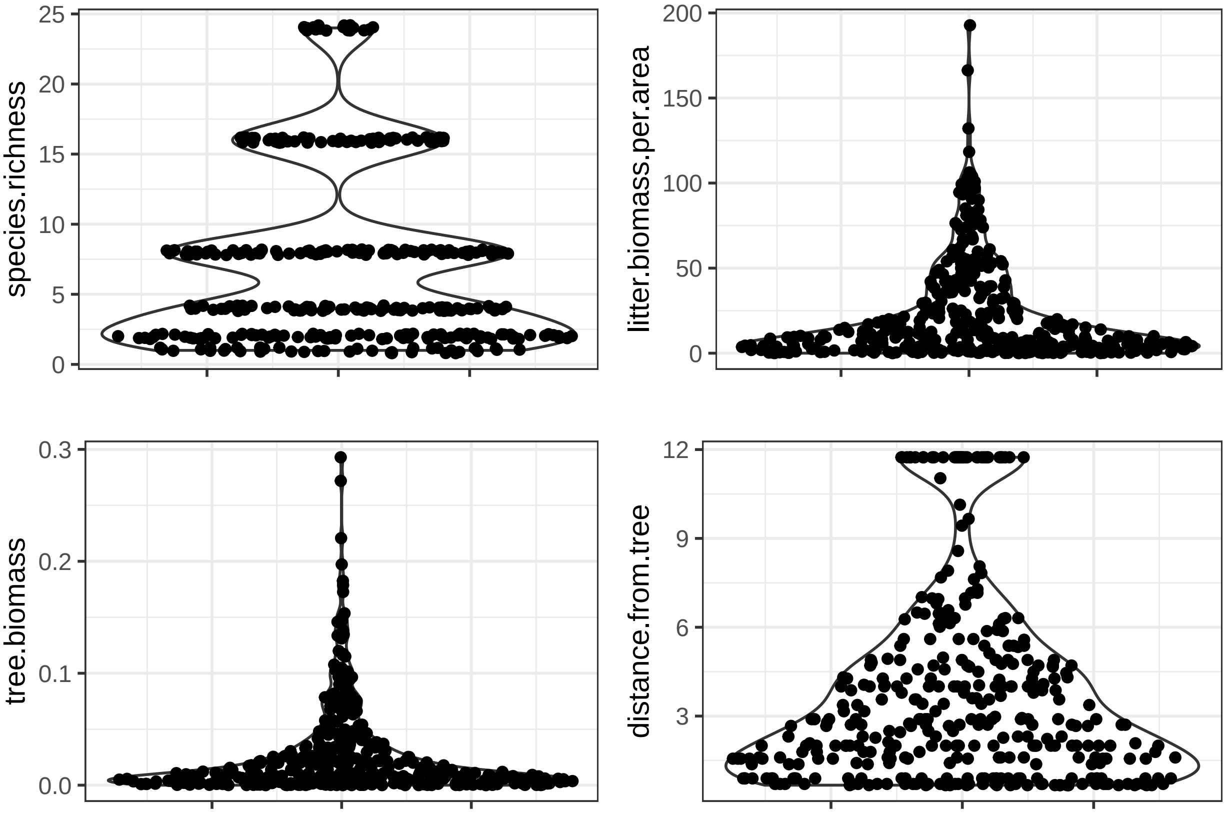

#### Model structure

```
stan_code = '
data{
  int n; // observations per species
  int m; // species number
  matrix[m,n] litter; // Species specific litter
  matrix[m,n] biomass;  // Species specific biomass
  matrix[m,n] dist;  // Species specific distance
  matrix[m,n] biodist;  // Species specific distance
}

parameters{
  vector<lower=0>[m] bio; // Species specific biomass parameter
  vector<lower=0>[m] d; // Species specific distance parameter
  vector<lower=0>[m] db;  // Species specific interraction parameter
  vector<lower=0>[m] sigma;  // Species specific sigma
}

model{
  // Priors
  matrix[m,n] mu;  // Species specific mu
  bio ~ normal(0,10); // Species specific biomass parameter prior
  d ~ normal(0,10);  // Species specific distance parameter
  db ~ normal(0,10);  // Species specific interaction parameter
  sigma ~ normal(0,10); // Species specific sigma

  // Likelihood
  for(i in 1:n){
    for(j in 1:m){
      mu[j,i] = bio[j] * biomass[j,i] + d[j] * dist[j,i] + db[j] * biodist[j,i] ;
      //Species specific litterfall
      litter[j,i] ~ normal(mu[j,i], sigma[j]);
    }
  }
}
'
```

```
## Running /Library/Frameworks/R.framework/Resources/bin/R CMD SHLIB foo.c
## using C compiler: 'Apple clang version 15.0.0 (clang-1500.0.40.1)'
## using SDK: 'MacOSX14.0.sdk'
## clang -arch arm64 -I"/Library/Frameworks/R.framework/Resources/include" -DNDEBUG   -I"/Library/Frame
## In file included from <built-in>:1:
## In file included from /Library/Frameworks/R.framework/Versions/4.3-arm64/Resources/library/StanHeade
## In file included from /Library/Frameworks/R.framework/Versions/4.3-arm64/Resources/library/RcppEigen/
## In file included from /Library/Frameworks/R.framework/Versions/4.3-arm64/Resources/library/RcppEigen/
## /Library/Frameworks/R.framework/Versions/4.3-arm64/Resources/library/RcppEigen/include/Eigen/src/Core
## namespace Eigen {
## ^
## /Library/Frameworks/R.framework/Versions/4.3-arm64/Resources/library/RcppEigen/include/Eigen/src/Core
## namespace Eigen {
##                ^
##                ;
## In file included from <built-in>:1:
```

```
## In file included from /Library/Frameworks/R.framework/Versions/4.3-arm64/Resources/library/StanHeade
## In file included from /Library/Frameworks/R.framework/Versions/4.3-arm64/Resources/library/RcppEigen/
## /Library/Frameworks/R.framework/Versions/4.3-arm64/Resources/library/RcppEigen/include/Eigen/Core:96
## #include <complex>
##          ^~~~~~~~~
## 3 errors generated.
## make: *** [foo.o] Error 1
```

#### Model fit

##### Posterior distribution

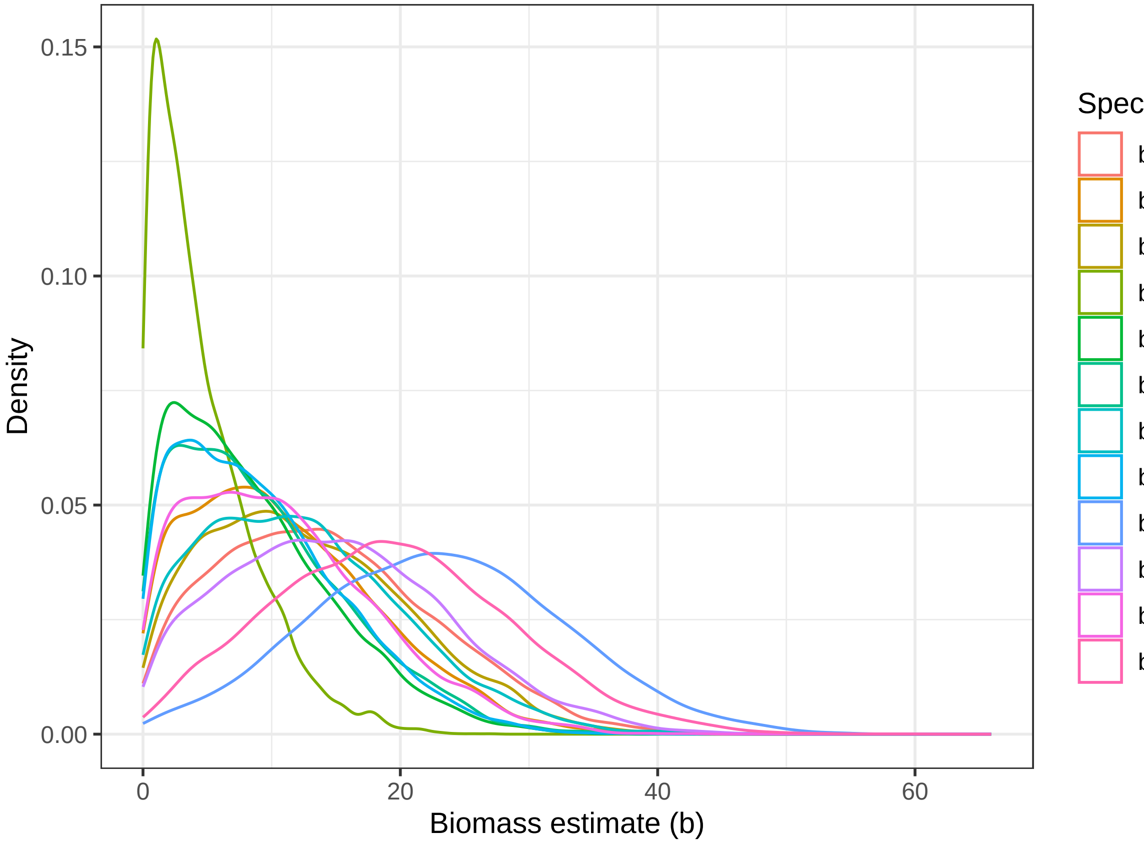


##### Biomass parameters

Average estimate:

```
##    bio[1]    bio[2]    bio[3]    bio[4]    bio[5]    bio[6]    bio[7]    bio[8]
## 14.298800  8.535652  9.280875 12.428158  9.222694 22.968842 15.195983 10.818468
##    bio[9]   bio[10]   bio[11]   bio[12]
## 19.342543 11.067364 12.819178  4.622921
```

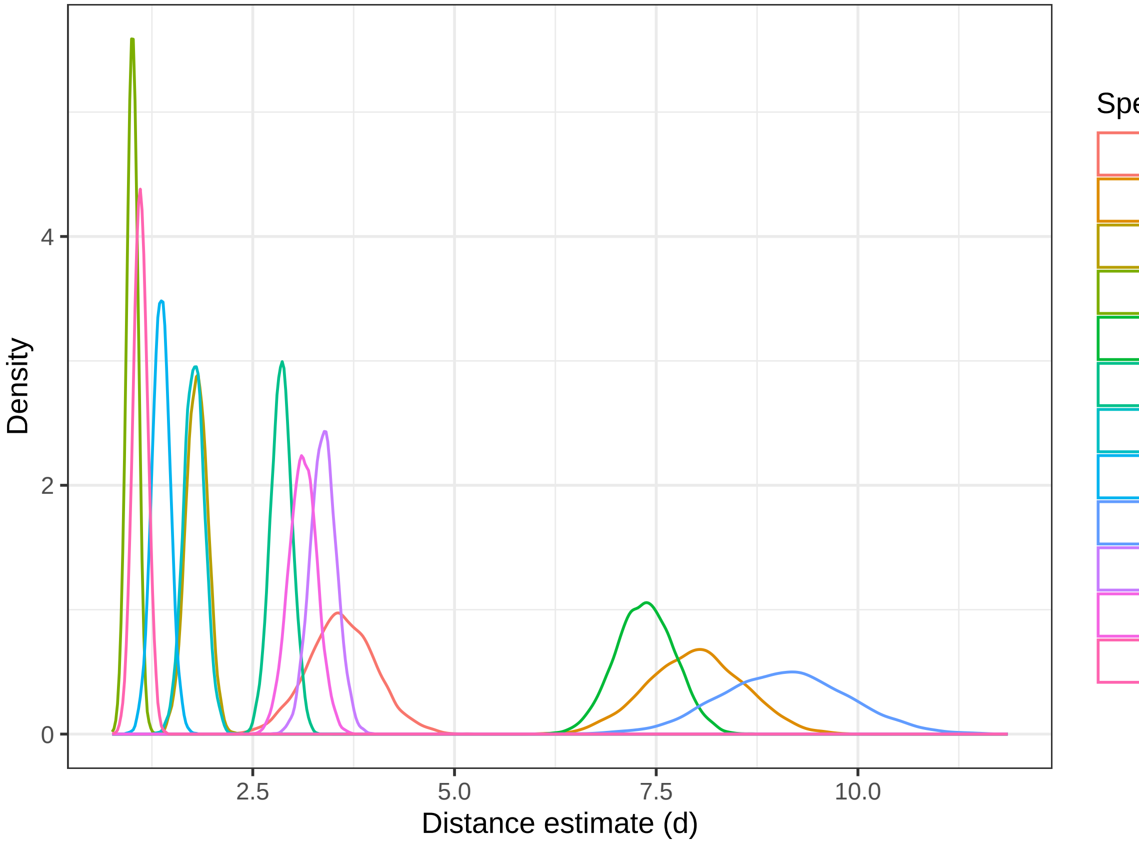


**Distance parameters**

Average estimate:

```
##     d[1]     d[2]     d[3]     d[4]     d[5]     d[6]     d[7]     d[8]
## 3.602818 7.363234 2.854709 1.774472 1.362265 9.085318 3.374269 3.120608
##     d[9]    d[10]    d[11]    d[12]
## 1.106374 7.990473 1.806871 1.007233
```

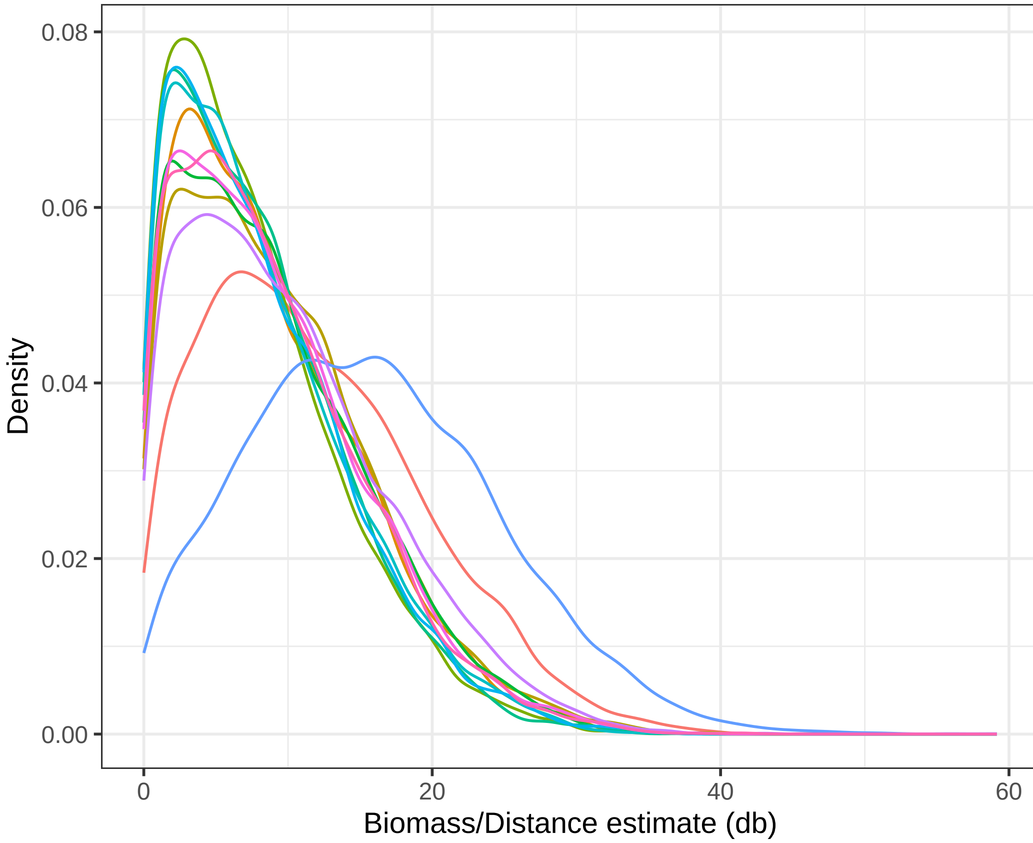


**Biomass-distance interaction**

Average estimate:

```
##     db[1]     db[2]     db[3]     db[4]     db[5]     db[6]     db[7]     db[8]
## 11.781653  8.967442  8.000781  8.207925  8.126577 15.870248  9.903248  8.914619
##     db[9]    db[10]    db[11]    db[12]
##  8.791810  8.819342  9.368846  7.776672
```

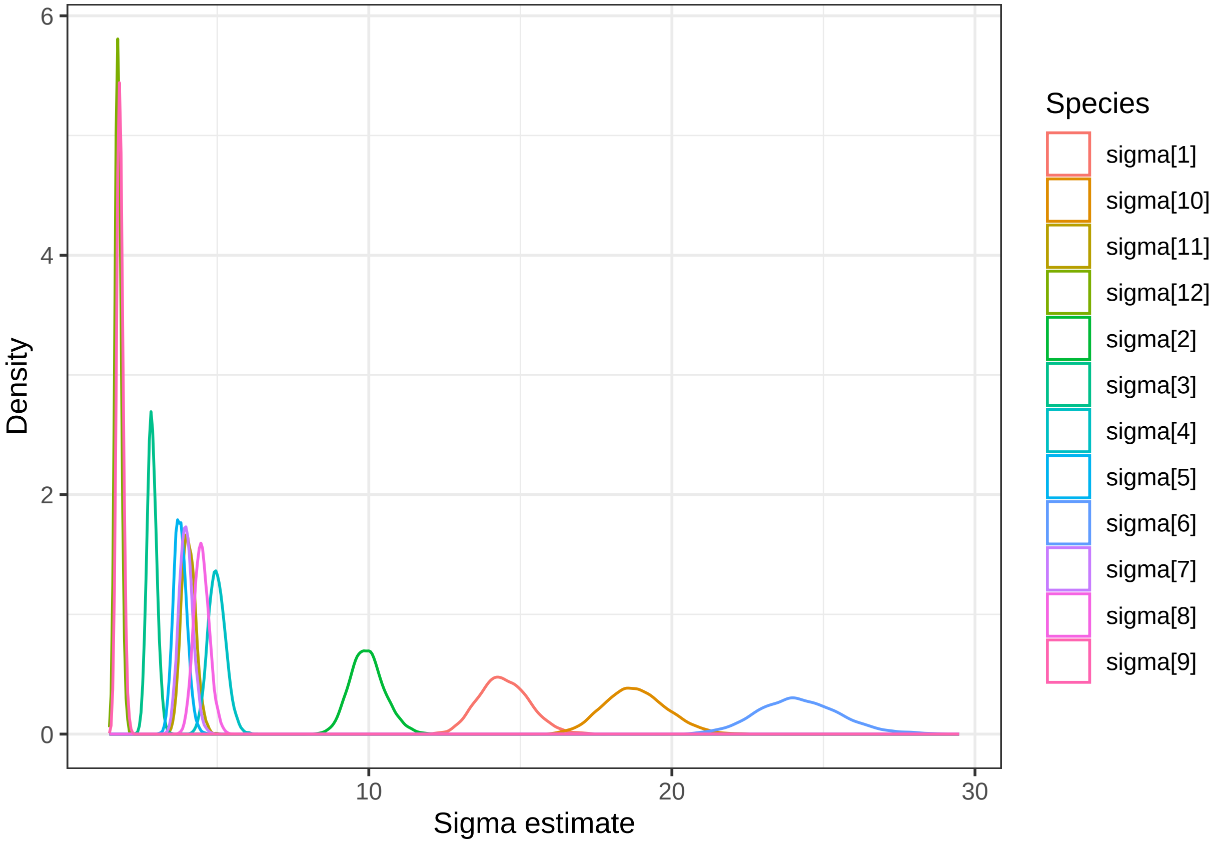


**Sigma**

**Species-specific R2**

```
##  [1] 0.3778935 0.7338101 0.7447753 0.6442434 0.5225354 0.5197896 0.7584071
##  [8] 0.7316106 0.5654657 0.5113701 0.5738229 0.5929612
```

```
## [1] 0.5628146
```

**LOO value**

```
## Warning: Relative effective sample sizes ('r_eff' argument) not specified.
## For models fit with MCMC, the reported PSIS effective sample sizes and
## MCSE estimates will be over-optimistic.
```

```
## Warning: Some Pareto k diagnostic values are too high. See help('pareto-k-diagnostic') for details.
```

```
##
## Computed from 8000 by 1920 log-likelihood matrix
##
##          Estimate    SE
## elpd_loo  -6669.3 351.7
## p_loo       880.2 312.6
## looic     13338.6 703.5
## ------
## Monte Carlo SE of elpd_loo is NA.
##
## Pareto k diagnostic values:
##                          Count Pct.    Min. n_eff
## (-Inf, 0.5]   (good)     1865  97.1%   1008
```

```
##  (0.5, 0.7]   (ok)          13   0.7%   126
##    (0.7, 1]   (bad)         12   0.6%   25
##    (1, Inf)   (very bad)    30   1.6%   1
## See help('pareto-k-diagnostic') for details.
```

#### Model predictions

```
## Warning in left_join(left_join(d.obs, d.predi, by = c("TSP", "name")), df, : Detected an unexpected m
## i Row 13 of `x` matches multiple rows in `y`.
## i Row 4 of `y` matches multiple rows in `x`.
## i If a many-to-many relationship is expected, set `relationship =
##   "many-to-many"` to silence this warning.
```

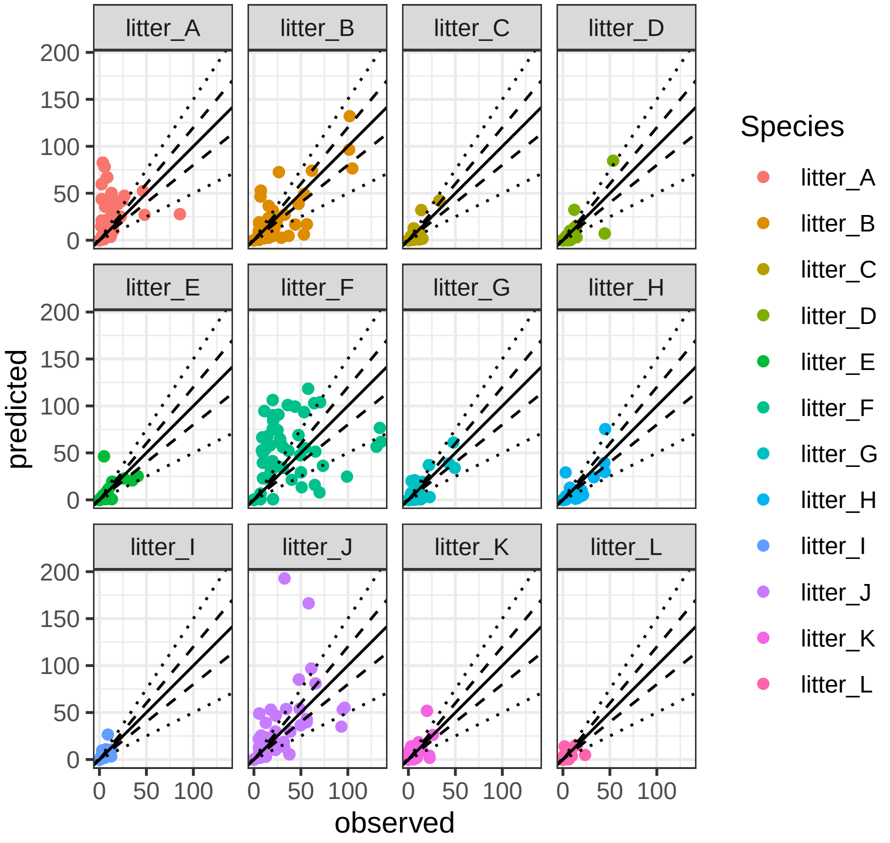

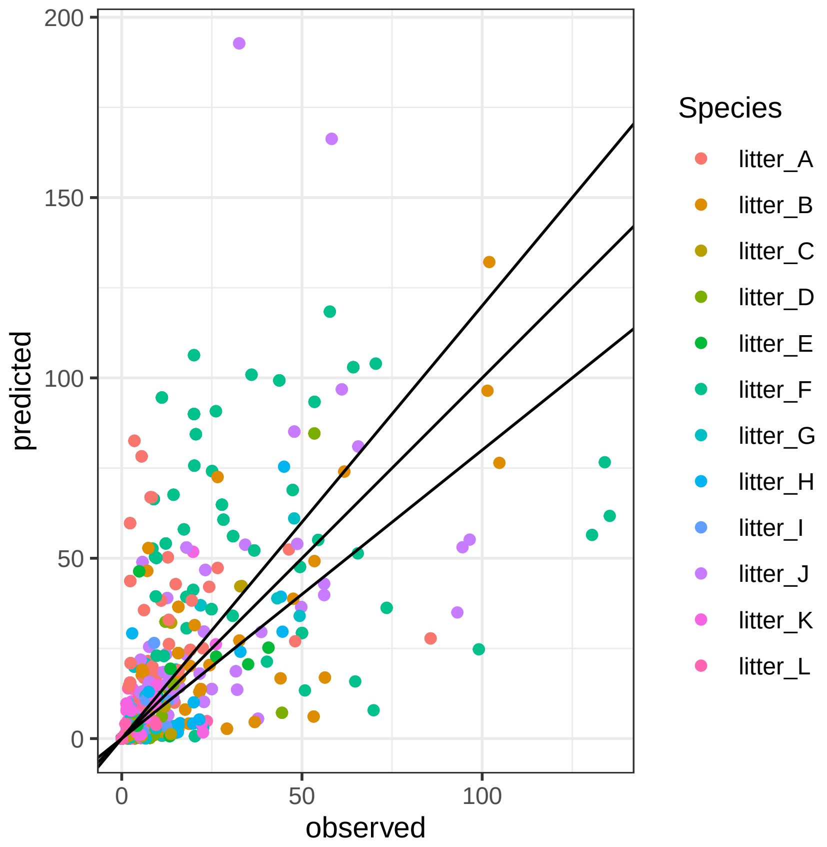
predicted
observed
0
50
100
150
200
Species
litter_A
litter_B
litter_C
litter_D
litter_E
litter_F
litter_G
litter_H
litter_I
litter_J
litter_K
litter_L

### Decomposition models

#### Datasets

##### Input data

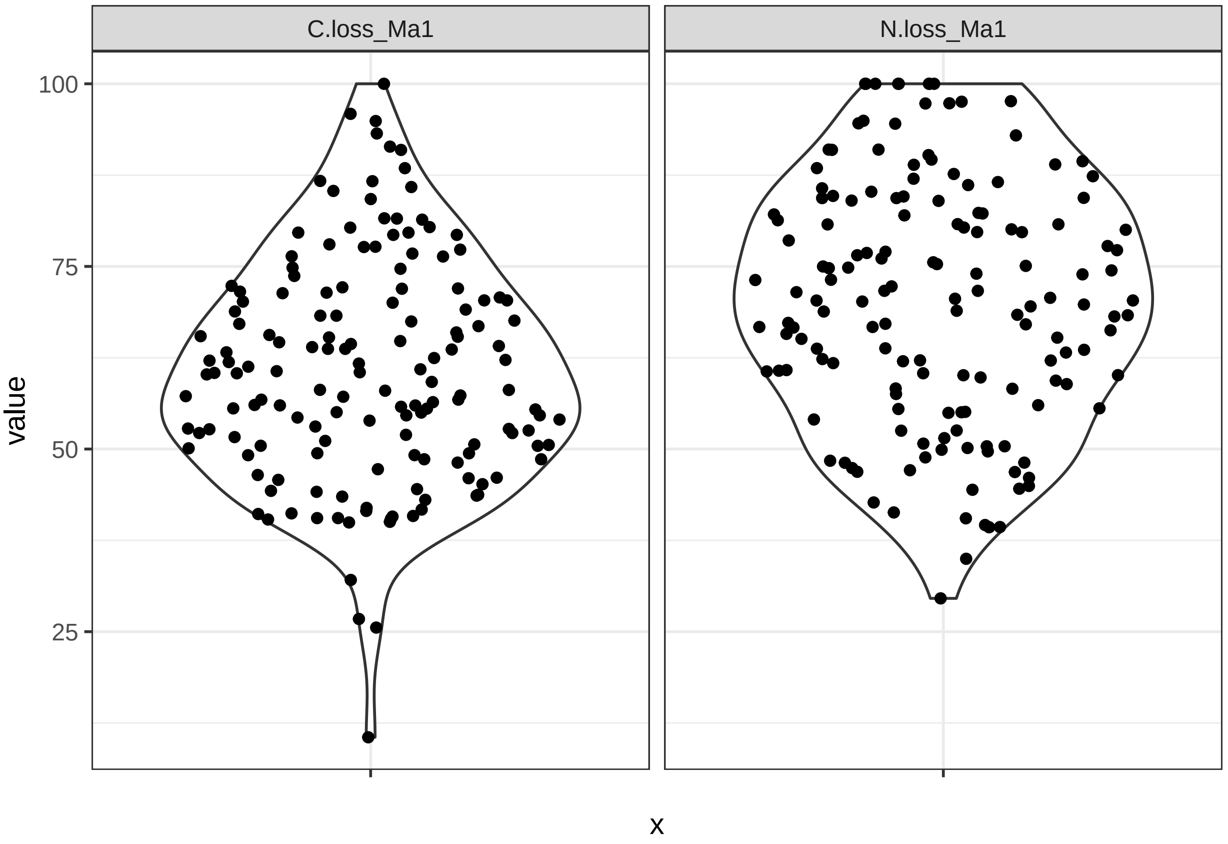


Species abundance

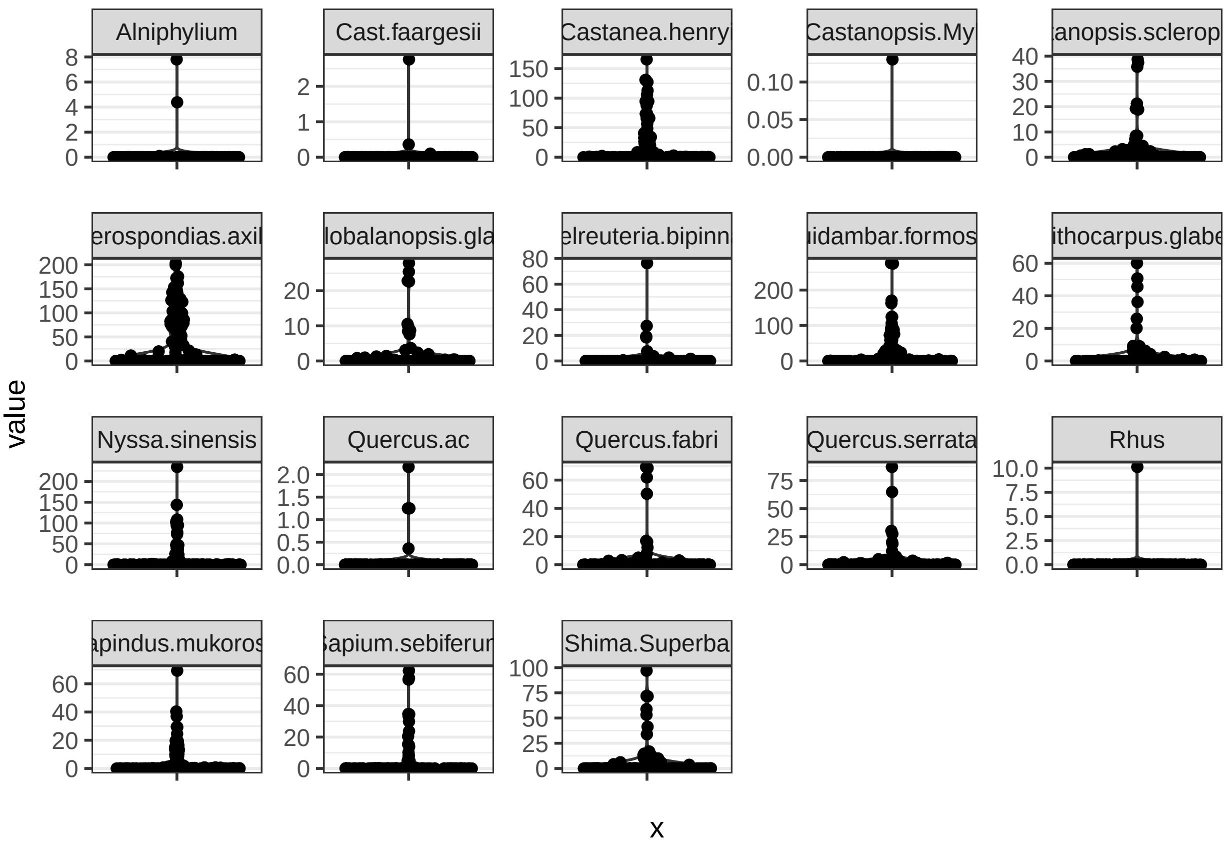


#### Model

```
stan_code = '
data{
  int n; // observations
  int m; // species number
  vector[n] decompC; // carbon decomposition
  vector[n] litterBM; // total litter biomass
  vector[n] litterSR; // litter species richness
  matrix[n, m] p; // proportion of species in litter biomass
}

parameters{
  vector[m] b; // species specific proportion parameter
  vector[m] a; // species specific interraction parameter
  real c; // litter biomass parameter
  real d; // litter species richness parameter
  real<lower=0> sigma; // sigma
}

model{
  // auxiliary variables
  real speff;
  real intereff;
```

```
  // Priors
  b ~ normal(0,10); // species specific proportion parameter
  a ~ normal(0,10); // species specific interraction parameter
  c ~ normal(0,10); // litter biomass parameter
  d ~ normal(0,10); // litter species richness parameter
  sigma ~ normal(0,10); // sigma

  // Likelihood

  for(k in 1:n){
    speff = 0.0;
    intereff = 0.0;
    for(i in 1:m){
      // summed species effects
      speff = speff + b[i] * p[k,i];
      // summed interaction effects: use trick for avoiding double sums
      intereff = intereff + a[i]*p[k,i]*(1-p[k,i]);
     }
    decompC[k] ~ normal(speff + intereff + c * litterBM[k] + d * litterSR[k], sigma);
  }
}
'
stan_model <- stan_model(model_code = stan_code)
```

```
## Trying to compile a simple C file
```

```
## Running /Library/Frameworks/R.framework/Resources/bin/R CMD SHLIB foo.c
## using C compiler: 'Apple clang version 15.0.0 (clang-1500.0.40.1)'
## using SDK: 'MacOSX14.0.sdk'
## clang -arch arm64 -I"/Library/Frameworks/R.framework/Resources/include" -DNDEBUG   -I"/Library/Framew
## In file included from <built-in>:1:
## In file included from /Library/Frameworks/R.framework/Versions/4.3-arm64/Resources/library/StanHeader
## In file included from /Library/Frameworks/R.framework/Versions/4.3-arm64/Resources/library/RcppEigen/
## In file included from /Library/Frameworks/R.framework/Versions/4.3-arm64/Resources/library/RcppEigen/
## /Library/Frameworks/R.framework/Versions/4.3-arm64/Resources/library/RcppEigen/include/Eigen/src/Core
## namespace Eigen {
## ^
## /Library/Frameworks/R.framework/Versions/4.3-arm64/Resources/library/RcppEigen/include/Eigen/src/Core
## namespace Eigen {
##                ^
##                ;
## In file included from <built-in>:1:
## In file included from /Library/Frameworks/R.framework/Versions/4.3-arm64/Resources/library/StanHeader
## In file included from /Library/Frameworks/R.framework/Versions/4.3-arm64/Resources/library/RcppEigen/
## /Library/Frameworks/R.framework/Versions/4.3-arm64/Resources/library/RcppEigen/include/Eigen/Core:96
## #include <complex>
##          ^~~~~~~~~
## 3 errors generated.
## make: *** [foo.o] Error 1
```

#### Model fit

```
# Carbon
dat_stan_C.Ma <- list(
```

```r
  n = nrow(df_hetero_simul),
  m = 16,
  decompC = df_hetero_simul$C.loss_Ma1,
  litterBM = rowSums(df_hetero_simul[,8:23]),
  litterSR = rowSums(df_hetero_simul[,8:23]!=0),
  p = df_hetero_simul[,8:23]/rowSums(df_hetero_simul[,8:24])
)

fit_carbon <- sampling(stan_model,
                       data = dat_stan_C.Ma,
                       iter = 3000,
                       warmup = 1000,
                       chains = 3,
                       cores = 3,
                       refresh=10,
                       control=list(max_treedepth=10)
)

# Nitrogen
dat_stan_N.Ma <- list(
  n = nrow(df_hetero_simul),
  m = 16,
  decompC = df_hetero_simul$N.loss_Ma1,
  litterBM = rowSums(df_hetero_simul[,8:23]),
  litterSR = rowSums(df_hetero_simul[,8:23]!=0),
  p = df_hetero_simul[,8:23]/rowSums(df_hetero_simul[,8:24]) # each row = 1 observations
)

fit_nitrogen <- sampling(stan_model,
                       data = dat_stan_N.Ma,
                       iter = 3000,
                       warmup = 1000,
                       chains = 3,
                       cores = 3,
                       refresh=10,
                       control=list(max_treedepth=10)
)
```

##### Model R2

Carbon

```
## [1] 0.490514
```

Nitrogen

```
## New names:
## * `` -> `...17`
## * `` -> `...18`
## * `` -> `...19`
```

```
## [1] 0.5361795
```

**Posterior distribution**

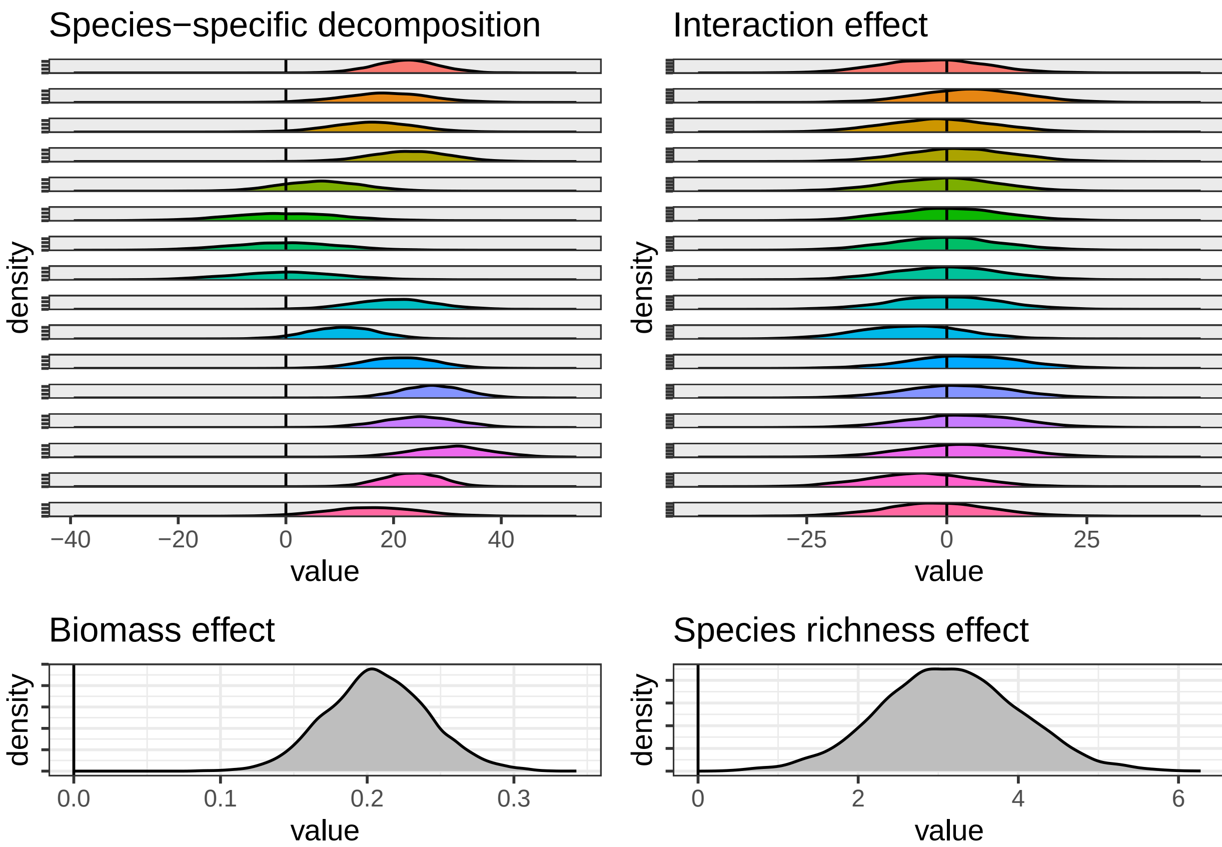


**Carbon model**

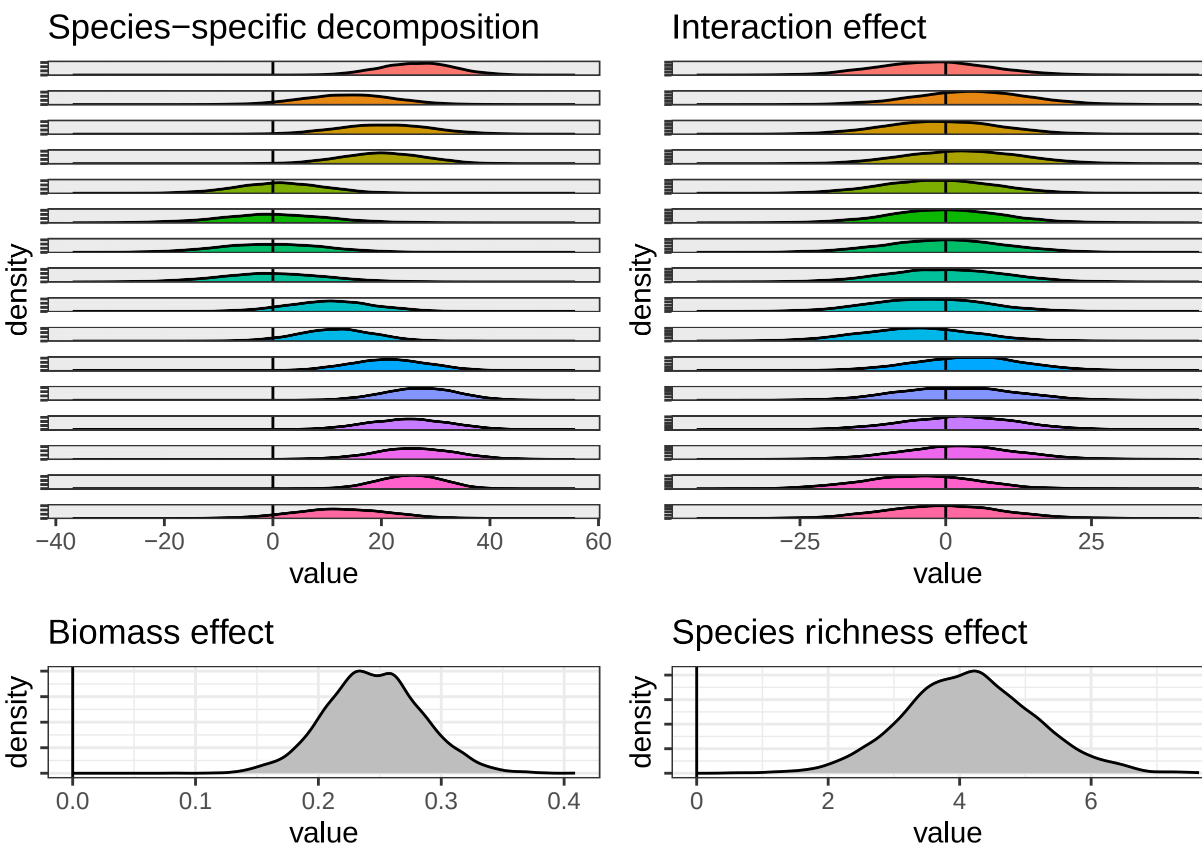


**Nitrogen model**

#### Model predictions

Carbon

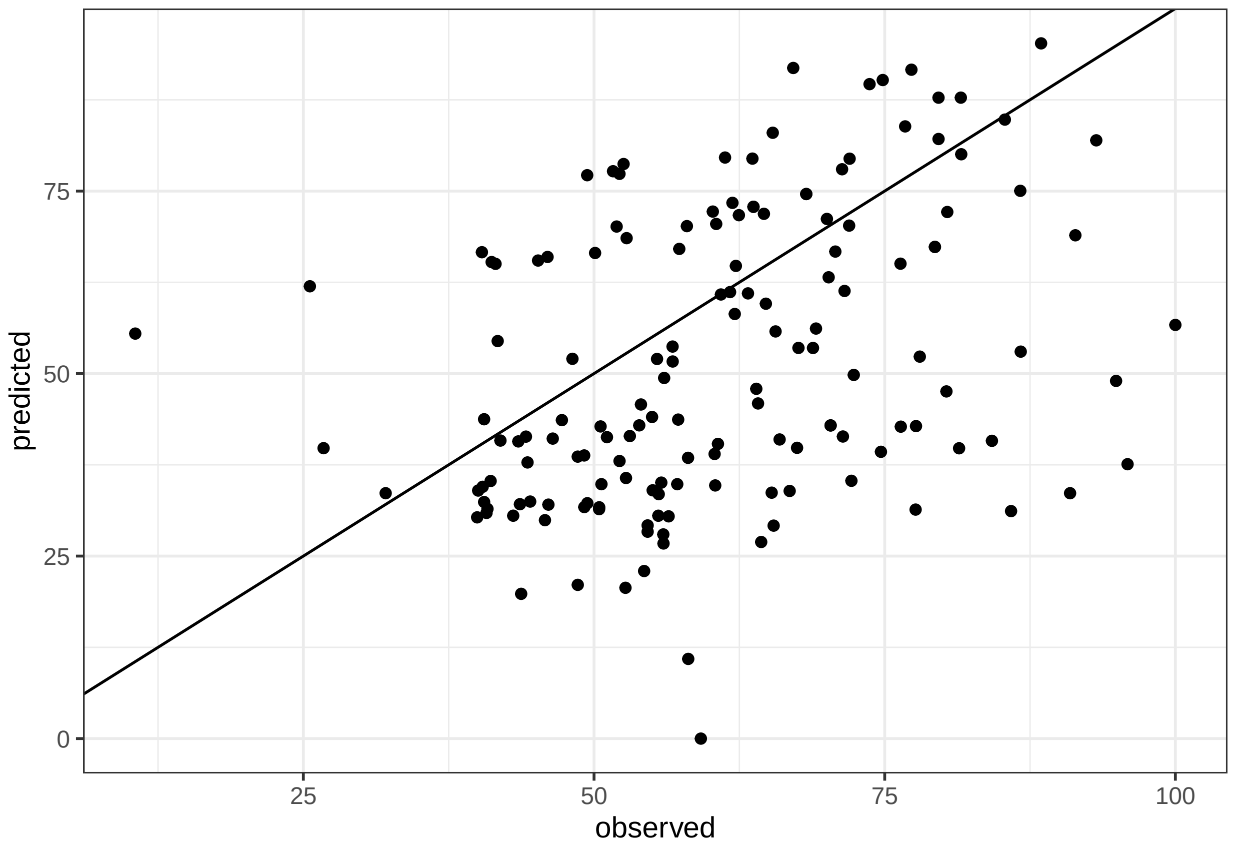
predicted
0
25
50
75
25
50
75
100
observed

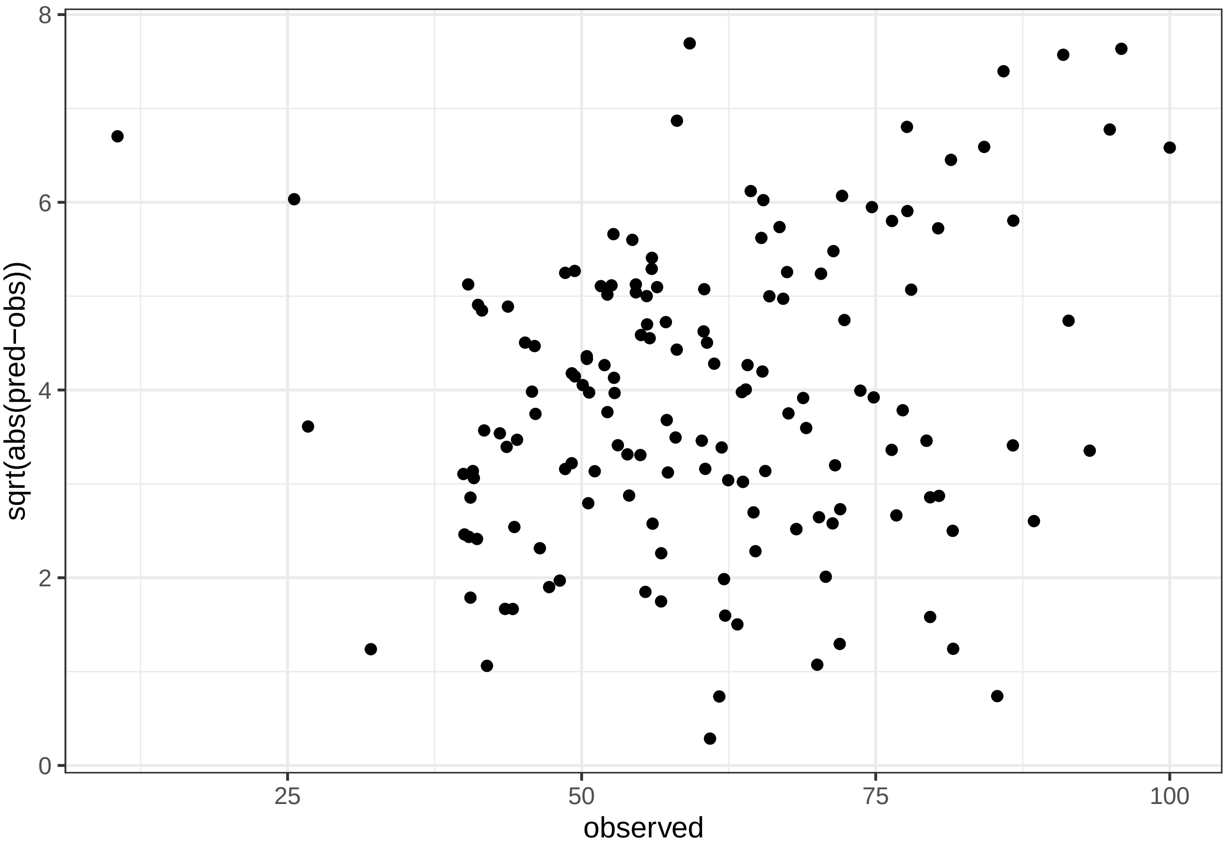


Nitrogen

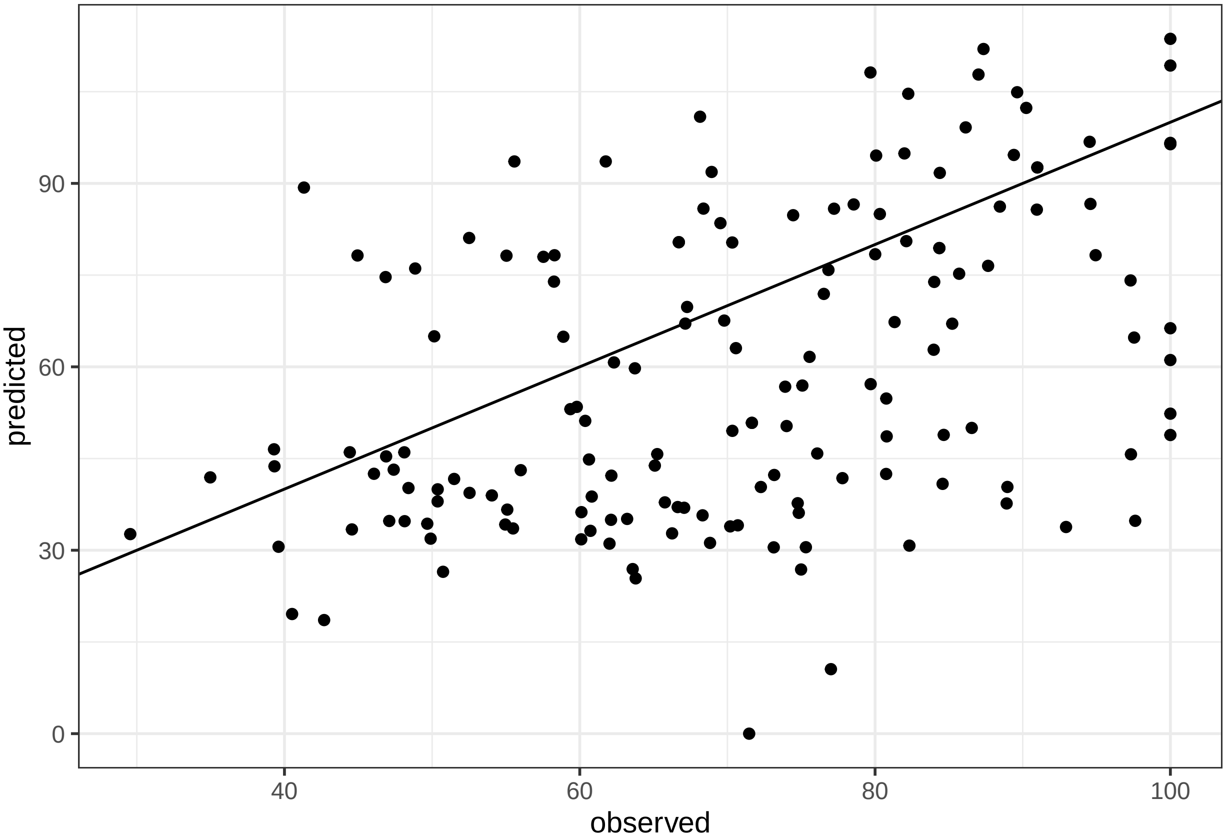

predicted
observed
0
30
60
90
40
60
80
100

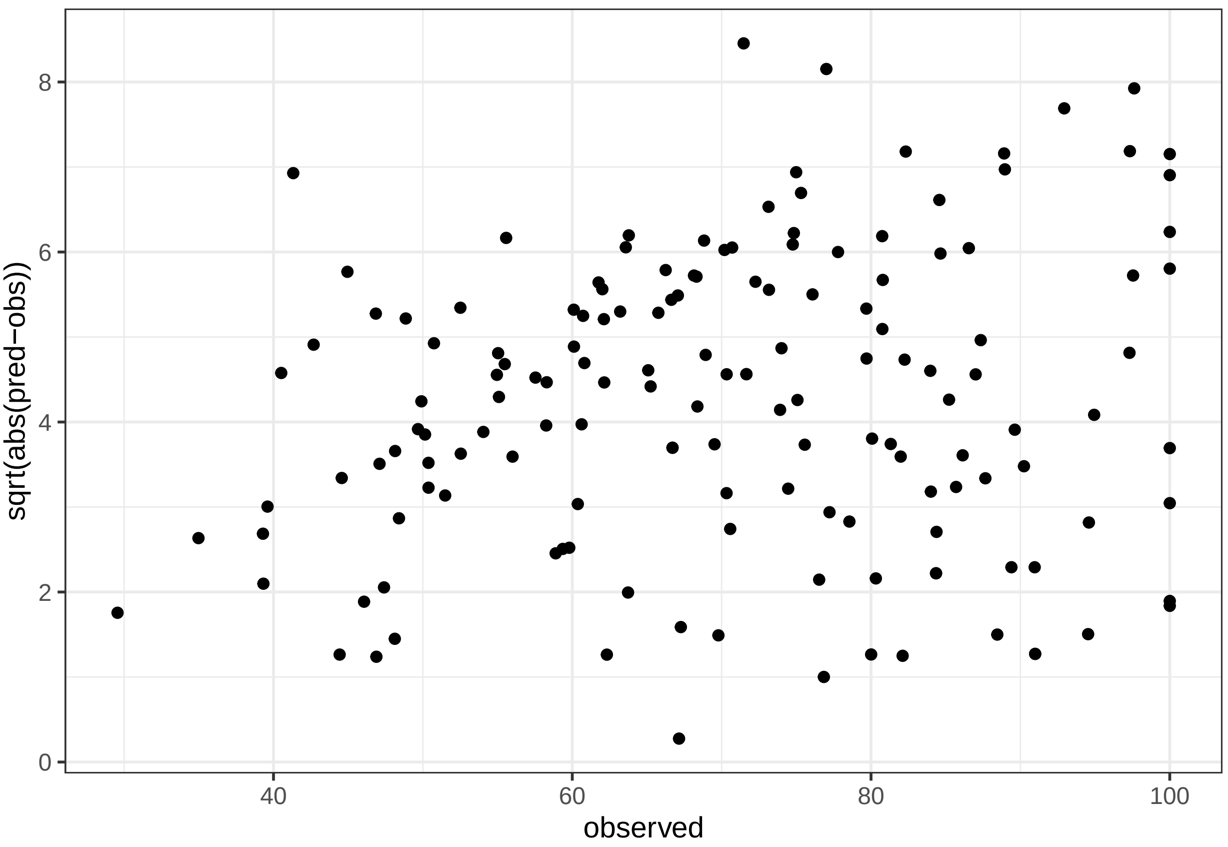

sqrt(abs(pred−obs))
0
2
4
6
8
40
60
80
100
observed
