## Supplementary material for "Improving forest ecosystem functions by optimizing tree species spatial arrangement": Suppl. S3

### Supplementary material S3

**9 species mixutes simulation outputs - additional variables**

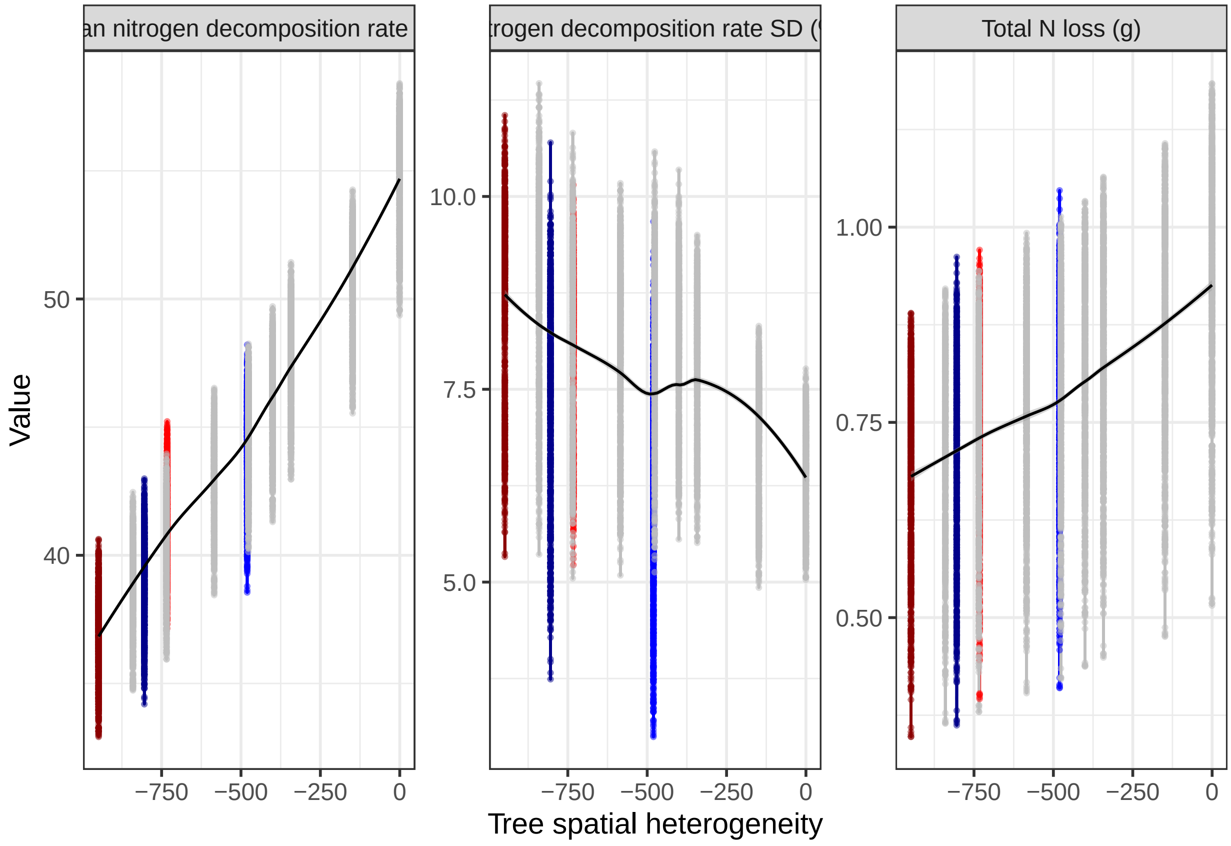

#### Across diversity outputs

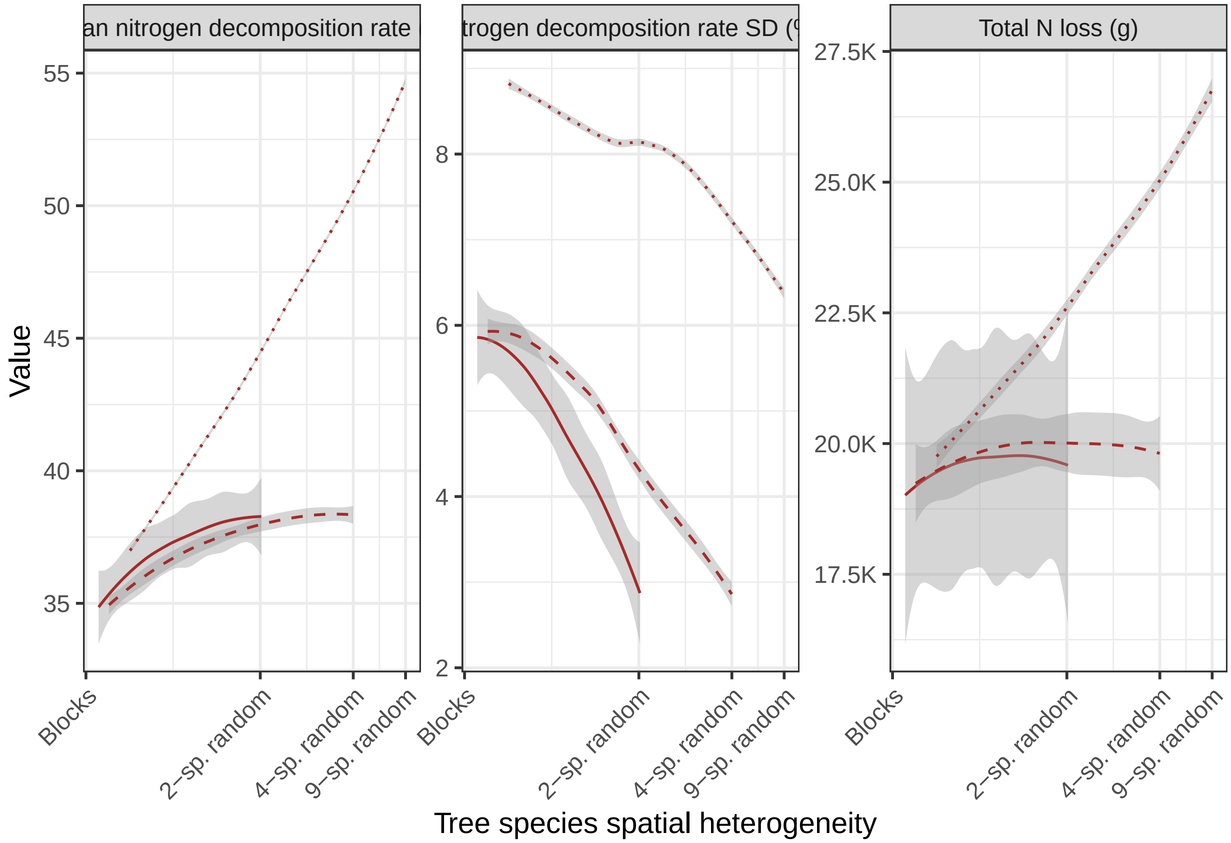

**Leaf traits**

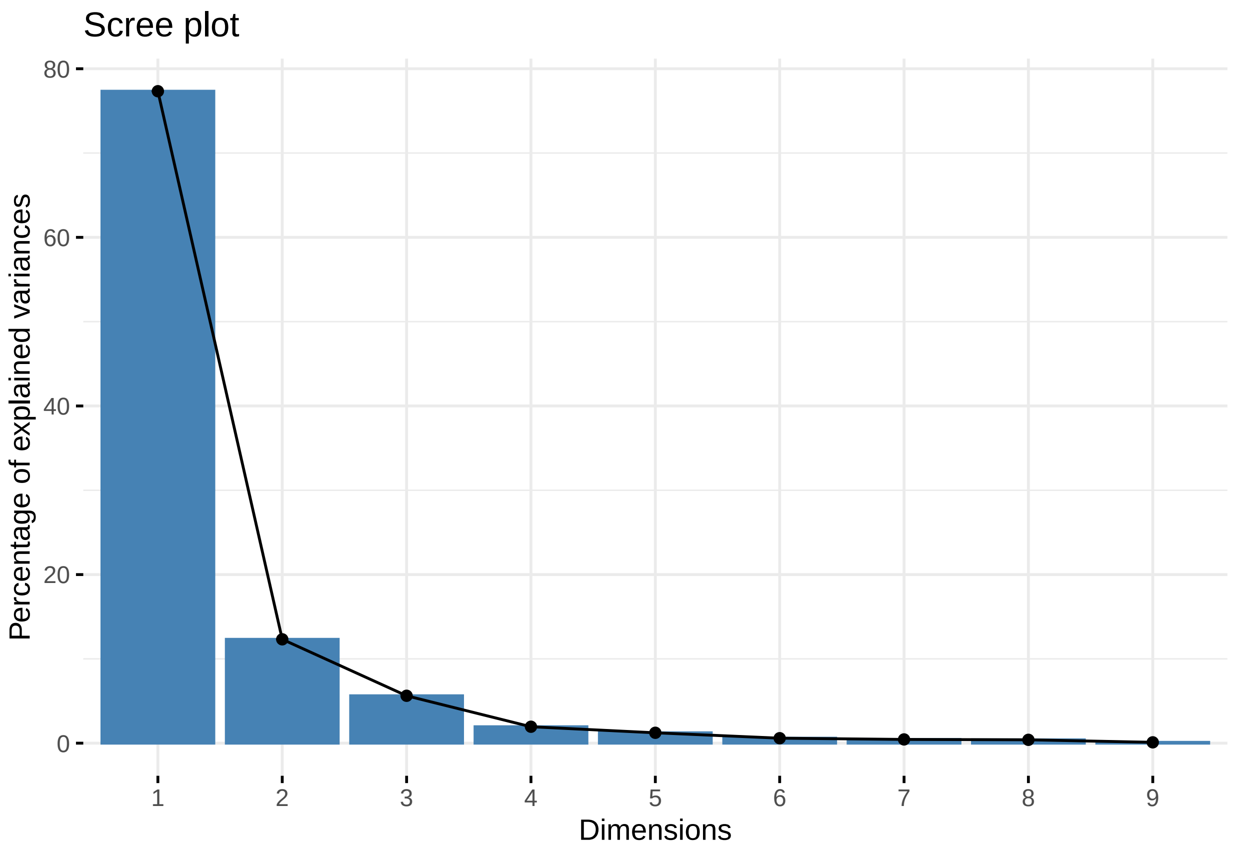
Scree plot
Percentage of explained variances
0
20
40
60
80
1
2
3
4
5
6
7
8
9
Dimensions

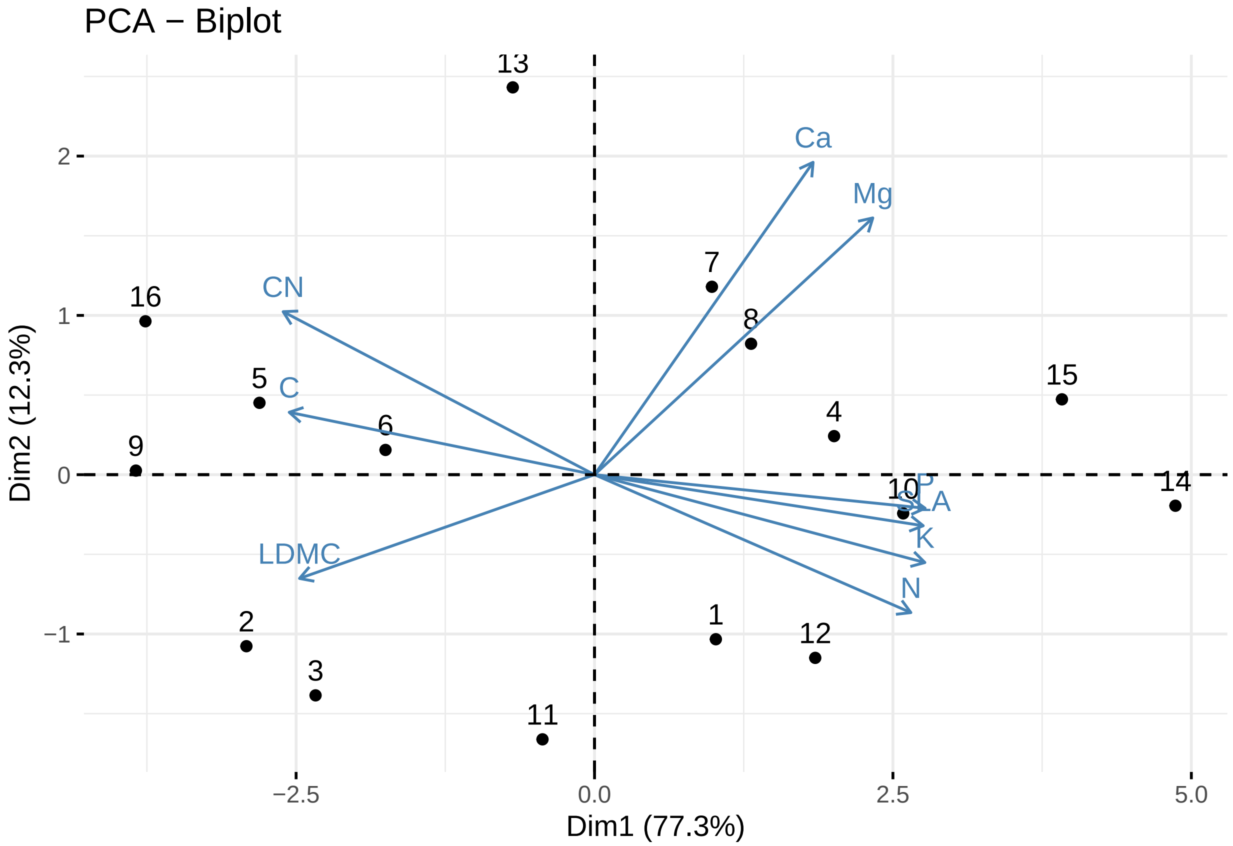
PCA − Biplot
Dim2 (12.3%)
Dim1 (77.3%)
−2.5
0.0
2.5
5.0
−1
0
1
2
Ca
Mg
CN
C
LDMC
P
SLA
K
N

#### Random forest partial plots

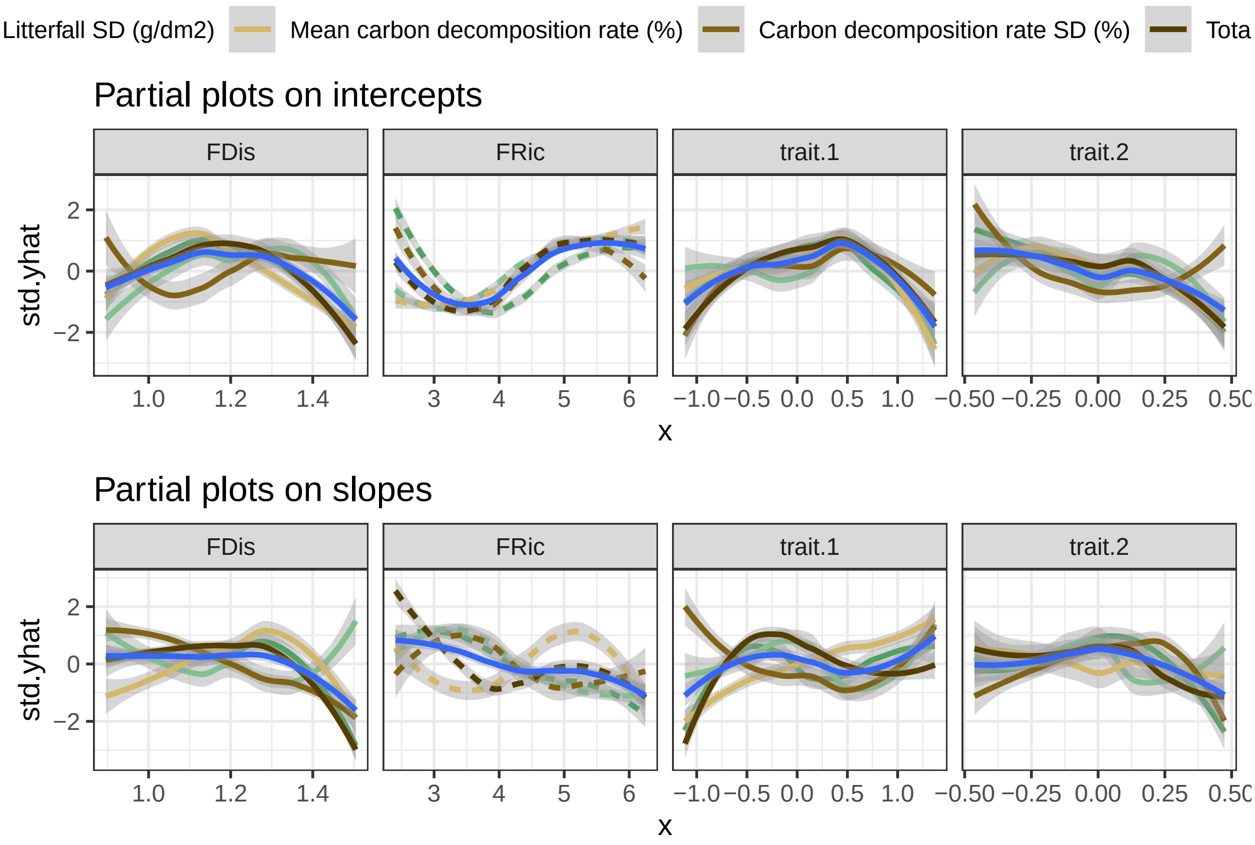
